# Circadian and Sex-Specific Responses to *Alternaria alternata* in Allergic Airway Inflammation

**DOI:** 10.64898/2026.09.30.755795

**Authors:** Santhosh Kumar Duraisamy, Joanna Mainzer, Chandrashekar Prasad, Steven Haworth, Mario Castro, Isaac Kirubakaran Sundar

## Abstract

Asthma severity and immune responses are regulated by circadian rhythms, yet the role of circadian timing in fungal allergen-induced airway inflammation remains poorly defined. *Alternaria alternata*, a clinically relevant fungal allergen, is strongly associated with severe asthma exacerbations and persistent airway dysfunction. Using a preclinical mouse model, we investigated whether pulmonary responses to *Alternaria* exposure are influenced by time of day and sex. We demonstrate that *Alternaria* induces allergic airway inflammation, airway hyperresponsiveness (AHR), and epithelial barrier disruption in a circadian- and sex-dependent manner. Female mice exhibited enhanced eosinophilic, neutrophilic, interstitial macrophage, and dendritic cell infiltration predominantly at Zeitgeber Time 0 (ZT0), whereas males showed stronger inflammatory responses at ZT12. *Alternaria* exposure increased IL-33 release and promoted ILC2 and Th2 cell accumulation, with distinct temporal patterns between sexes. Cytokine and chemokine profiling revealed robust Th2, proinflammatory, and Th17 responses that varied with exposure time, with higher responses at ZT12 in females and ZT0 in males. Markers of tissue injury, including lactate dehydrogenase and uric acid, followed similar circadian-dependent trends. Functionally, *Alternaria*-exposed mice displayed greater AHR during the resting phase, particularly in males. Additionally, *Alternaria* disrupted epithelial junctional proteins (E-cadherin and ZO-1) and altered expression of core circadian clock proteins CLOCK, BMAL1, and REV-ERBα. Mechanistically, peak inflammation coincided with enhanced STAT1, STAT3, and STAT6 activation. Collectively, these findings identify circadian timing and sex as key determinants of fungal allergen-induced airway inflammation and suggest chronotherapeutic approaches may improve asthma outcomes.

## Introduction

Physiological processes in vertebrates, including feeding behavior, metabolism, inflammation, and the sleep–wake cycle, are tightly regulated by the circadian clock ^1^. In mammals, nearly all cells exhibit intrinsic diurnal oscillations coordinated by interactions between the central and peripheral clocks. The master pacemaker, located in the suprachiasmatic nucleus (SCN), synchronizes peripheral rhythms through neuronal and hormonal cues in response to environmental signals such as light ^1^. At the molecular level, circadian rhythms arise from transcription–translation feedback loops involving core clock components such as BMAL1, CLOCK, PER1/2, CRY1/2, and the nuclear receptors REV-ERBα/β ^2^. In the lung, airway epithelial cells, smooth muscle cells, and immune cells harbor functional circadian oscillators that regulate gene programs controlling inflammation, epithelial barrier integrity, and airway tone ^3^.

Disruption of circadian rhythms perturb physiological homeostasis and is associated with increased morbidity and mortality ^4, 5^. Asthma exhibits strong circadian variation, with symptoms, airway obstruction, and risk of fatal exacerbations peaking at night or in the early morning ^6^. Pulmonary function indices, including peak expiratory flow and FEV, also follow a diurnal pattern, reaching their lowest levels in the early morning ^7^. Dysregulated circadian timing amplifies airway inflammation, enhances bronchial hyperresponsiveness, and contributes to increased asthma severity ^2^.

Fungal allergens are among the strongest environmental drivers of severe asthma ^8, 9^. *Alternaria alternata* is clinically significant due to its potent allergens, such as Alt a1, which elicit strong IgE-mediated responses ^10^. Epidemiological studies consistently link airborne Alternaria exposure to increased asthma symptoms, hospitalization, and life-threatening attacks ^11^, with sensitization especially prevalent among children and young adults with persistent airflow limitation and severe disease ^12^. Mechanistically, Alternaria secretes proteases that compromise epithelial barrier integrity and enhance allergen penetration ^13^. Alternaria exposure rapidly activates airway epithelial cells and triggers release of alarmins, including IL-33 and TSLP, which subsequently activate type 2 innate lymphoid cells (ILC2s) and initiate type 2 inflammation ^14, 15^. Notably, airborne Alternaria spores themselves exhibit diurnal oscillation influenced by humidity, temperature, and agricultural activity ^16^, suggesting that fungal exposure may intersect with circadian control of airway immunity.

Signal transducer and activator of transcription (STAT) proteins are essential mediators of cytokine signaling and immune regulation. Both STAT6 and STAT3 are critically involved in asthma pathogenesis ^17, 18^. STAT6 activation by IL-4 and IL-13 drives Th2 differentiation, IgE class switching, and mucus hypersecretion, the hallmarks of allergic asthma ^19, 20^. In contrast, STAT3 activation by IL-6 promotes Th17 differentiation, neutrophilic inflammation, and airway remodeling, features associated with severe and corticosteroid-refractory asthma ^21^. Alternaria exposure modulates STAT-dependent pathways. STAT6-driven epithelial FIZZ1 expression promotes IgE production, eosinophilia, mucus hypersecretion, and airway thickening ^22^, while Alternaria-induced epithelial injury and alarmin release (IL-33, IL-21, IL-23, IL-6) may activate STAT3-dependent pathways, contributing to Th17 responses, neutrophilic inflammation, and airway remodeling ^23, 24^. Dysregulated STAT signaling may therefore represent a mechanistic link between fungal sensitization and severe asthma phenotypes.

Despite increasing recognition that circadian timing influences immune responses, it remains unclear whether Alternaria exposure elicits time-of-day or sex-specific differences in inflammatory cell recruitment and STAT pathway activation in the lung. Moreover, the interaction between circadian clock components and allergen-driven inflammatory pathways remains poorly understood, particularly in the context of clinically relevant fungal allergens such as *Alternaria*.

To address these gaps, we employed comprehensive *in vivo* mouse models to investigate the time-of-day- and sex-dependent regulation of *Alternaria alternata*–induced allergic airway inflammation. By defining circadian gating of inflammatory and STAT-dependent pathways, this study provides new insight into the temporal regulation of allergen-driven airway responses and highlights the potential of chronotherapeutic strategies to mitigate asthma severity.

## Material and Methods

### Mice

Adult C57BL/6 (WT) mice, females and males (2–3 months old) were housed under a 12:12 h light–dark cycle in the Research Support Facility (RSF) vivarium at the University of Kansas Medical Center (KUMC). All procedures were approved by the Institutional Animal Care and Use Committee (IACUC) at KUMC (ACUP #2020-2575-1).

### Acute *Alternaria alternata* (ALT) asthma model

Adult C57BL/6 mice (2–3 months old; females and males) were exposed to an acute ALT-induced allergic airway inflammation model. *Alternaria alternata* extract (Greer Laboratories, Lenoir, NC) was administered intranasally once daily for 4 consecutive days under light isoflurane (5%) anesthesia. Mice received ALT dose (10 µg in 30 µL sterile PBS) or control (30 µL PBS) at circadian times: ZT0 (06:00; beginning of rest phase) or ZT12 (18:00; beginning of active phase). These Zeitgeber times were selected to evaluate circadian influences on ALT-induced airway inflammation. Mice were euthanized 24 hours after the final PBS or ALT exposure to assess acute allergic responses.

### Airway hyperresponsiveness (AHR) measurement

Airway hyperresponsiveness was assessed 24 hours after the final ALT exposure using increasing concentrations of methacholine (0–90 mg/mL) with the flexiVent system (SCIREQ Scientific Respiratory Equipment Inc., Montreal, Canada), as previously described ^25^. Briefly, mice were anesthetized, tracheostomized, and mechanically ventilated, followed by delivery of aerosolized methacholine in a dose-dependent manner.

Respiratory system resistance (Rrs), elastance (Ers), compliance (Crs), Newtonian resistance (Rn), and tissue damping (G) were calculated using flexiVent software. Mean dose-response curves for each parameter were generated after normalization to baseline values obtained prior to methacholine challenge, as described previously ^26^.

### Bronchoalveolar Lavage (BAL) Fluid

Mice were euthanized by intraperitoneal administration of Euthaphen (pentobarbital sodium; 50 mg/kg body weight), followed by exsanguination. BAL fluid was collected by gently lavaging the lungs three times with 0.6 mL sterile saline (0.9% NaCl) as previously described ^27^. The recovered lavage fluid was centrifuged at 2,000 rpm to separate cellular and acellular components. Cell-free BAL supernatants were aliquoted and stored at −80°C for subsequent analysis of inflammatory mediators. The BAL cell pellet was resuspended in 1 mL FACS staining buffer, and total cell counts were obtained using a Countess 3 Automated Cell Counter (Thermo Fisher Scientific, Waltham, MA, USA). Briefly, 10 µL of the resuspended cells was mixed with 10 µL of acridine orange/propidium iodide (AOPI) viability dye and loaded onto a Countess cell counting chamber slide. Live and dead cells were quantified according to the manufacturer’s protocol.

### Flow Cytometry Analysis

Cells obtained from BAL fluid (1 × 10 cells in 100 µl) were resuspended in FACS staining buffer and incubated with anti-CD16/32 (BioLegend) to block Fc receptors. Cells were then stained with the following antibodies. Myeloid panel: CD45 BV605, CD3 PE-Cy7, CD11b APC-Cy7, IA/IE BV650, CD11c BV785, CD24 AF488, CD193 (CCR3) PE, Gr-1 AF700, CD64 APC, and Siglec-F PE-CF594. Lymphoid panel: CD45 BV605, CD90.2 BV785, CD3 PE-Cy7, CD4 APC, CD44 AF700, and CD19 APC-Cy7. Intracellular transcription factors: T-bet BV710, GATA3 PE, and ROR-γt PE-ef610. Ghost Dye 510 (Tonbo Biosciences) was used to exclude dead cells. A detailed list of all antibodies, catalog numbers, and vendors is provided in **Table S1** (Key Resources). Flow cytometry data were acquired on a BD LSRFortessa by collecting 100,000 events per sample for myeloid analyses and 1,000,000 events for lymphoid subset analyses. Compensation was performed using OneComp eBeads (Thermo Fisher Scientific) along with single-color controls and unstained controls. Data was analyzed using FCS Express 7 software.

### Histological Analysis and Immunohistochemical Staining

The large left lung lobe was inflated with 1% low-melt agarose in 1× PBS, followed by fixation in 10% neutral buffered formalin for 48 hours. Post-fixation, tissues were progressively dehydrated in 30%, 50%, and 70% ethanol (20 minutes each) and subsequently embedded in paraffin. Midsagittal lung sections (5 μm) were cut using a rotary microtome for histopathological staining and semi-quantitative analysis.

### Hematoxylin and Eosin (H&E) Staining

H&E staining was performed to assess peribronchial (airway), alveolar, and perivascular inflammation. Lung sections were scored manually and in a blinded manner using a five-point grading scale (0–4). 0 – Normal lung; no appreciable inflammatory infiltrate; 1 – Minimal inflammation; few scattered immune cells without layering; 2 – Mild inflammation; several layers of inflammatory cells around airways or vessels but not dense enough to encircle adjacent structures; 3 – Moderate inflammation; four to five well-defined layers of immune cells, often dense enough to border nearby structures; 4 – Severe inflammation; extensive infiltration with > 4–5 layers of cells fully encircling the airway/vascular structures and infiltrating surrounding tissue. When histological features fell between two categories, an intermediate (averaged) score was assigned. The mean inflammation score for each animal was calculated and used for statistical comparisons between PBS control and ALT-exposed groups.

### Gene Expression by Quantitative Real-Time PCR Analysis

Lung tissues harvested from PBS control and ALT-exposed mice were flash-frozen in liquid nitrogen and stored at −80 °C. Lung tissues (∼50 mg) were homogenized in Trizol, and total RNA was isolated using the RNeasy mini kit (Qiagen) according to the manufacturer’s instructions. Total RNA quality and quantity were measured using a Nanodrop spectrophotometer (Nanodrop One; Thermo Scientific) followed by cDNA synthesis (1 μg of total RNA) with the First Strand cDNA synthesis kit (Qiagen). Prepared cDNA was stored at −20°C and used for qRT-PCR analysis using CFX 96 Real-Time PCR system (Bio-Rad). Gene expression was determined using quantification cycle (Cq) values, and relative expression was calculated using the 2^−ΔΔCt^ method, with *Rplp0* serving as the internal reference gene. The list of gene-specific qRT-PCR primers used in the study is provided (**Table S2**).

Due to space constraints, detailed descriptions of experimental procedures and data analysis approaches are provided in the **Supplementary Methods** section.

### Statistical Analysis

The statistical analysis was carried out using GraphPad Prism 9 software (La Jolla, CA). Differences between two or more experimental groups were assessed by Two-way ANOVA using Tukey’s multiple comparison test. Three-way interaction analysis between Treatment (PBS vs. ALT), Sex (Females vs. Males), and Time (ZT0 vs. ZT12) was performed using the generalized linear modeling through the R statistical computing language (**Table S3**). Results were presented as the mean ± SEM. A *P-*value less than 0.05 was considered statistically significant.

## Results

### Alternaria exposure induces time-of-day-dependent immune infiltration into BAL fluid

An acute *Alternaria alternata* (ALT)-induced allergic asthma model was used to determine whether airway exposure elicits time-of-day–dependent immune infiltration into the lung, comparing responses at ZT0 (6:00 am; resting phase) and ZT12 (6:00 pm; active phase) in female and male mice. ALT exposure significantly increased total bronchoalveolar lavage (BAL) cellularity at both ZT0 and ZT12 in females and males compared with their respective PBS controls. Notably, ALT-induced infiltration was significantly greater during the resting phase (ZT0) than the active phase (ZT12) in males only (**Figure 1A**).

**Figure 1.**
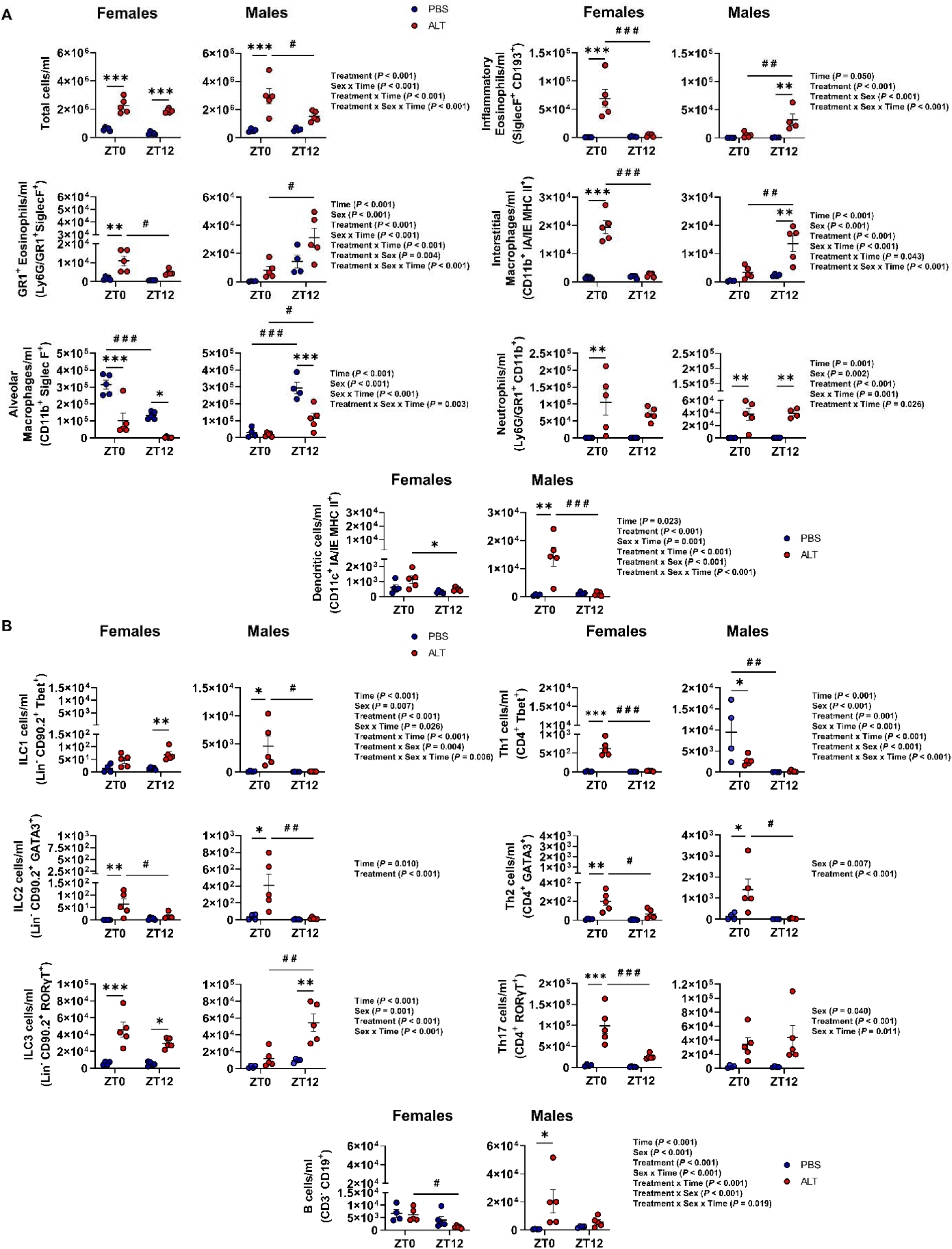
*Alternaria alternata* (ALT) exposure exhibits time-of-day and sex-dependent differences in immune cell infiltration into the lung. C57BL/6 wild-type female and male mice were administered ALT extract daily for four consecutive days. (A–B) ALT exposure differentially influenced infiltration of myeloid and lymphoid cell populations in PBS- and ALT-treated female and male mice. Immune cell infiltration following ALT challenge was quantified using multicolor flow cytometry. Data represent mean ± SEM (n = 4–5/group). Statistical analysis was performed using two-way ANOVA followed by Tukey’s multiple comparison test. * *P* < 0.05, ** *P* < 0.01, *** *P* < 0.001 compared with the respective PBS control at ZT0 or ZT12; ^#^*P* < 0.05, ^#^ ^#^*P* < 0.01, ^#^ ^#^ ^#^ *P* < 0.001 between ALT-exposed groups at ZT0 vs. ZT12. Interactions among Treatment x Sex x Time were analyzed using generalized linear modeling in R (see Supplemental Table S3).

Among infiltrating granulocytes and myeloid populations, ALT-exposed females exhibited significantly higher numbers of GR1 eosinophils (GR1 EOS), inflammatory eosinophils (iEOS), and interstitial macrophages (IMs) at ZT0 compared with ZT12, with neutrophils showing a similar, though not statistically significant, trend. In contrast, ALT-exposed males showed higher GR1 EOS, iEOS, and IMs at ZT12 relative to ZT0, with no evidence of time-of-day variation in neutrophil infiltration. Alveolar macrophages (AMs) were reduced following ALT exposure in both sexes at ZT0 and ZT12 compared with PBS controls.

Interestingly, baseline AM numbers exhibited sex-specific diurnal variation, with higher AM counts at ZT0 in PBS-treated females and at ZT12 in PBS-treated males. In addition, dendritic cells (DCs) were significantly increased at ZT0 compared with ZT12 following ALT exposure in males but slightly increased in females (**Figure 1A**).

Interaction analysis of BAL myeloid cell populations revealed significant main effects and interactions across several parameters. Total BAL cell counts showed significant effects of Treatment, Sex × Time, Treatment × Time, and Treatment × Sex × Time. Significant effects were also observed for iEOS (Time, Treatment, Treatment × Sex, Treatment × Sex × Time), GR1 EOS (Time, Sex, Treatment, Sex × Time, Treatment × Time, Treatment × Sex, Treatment × Sex × Time), IMs (Time, Sex, Treatment, Sex × Time, Treatment × Time, Treatment × Sex × Time), AMs (Time, Sex, Sex × Time, Treatment × Sex × Time), neutrophils (Time, Sex, Treatment, Sex × Time, Treatment × Time), and DCs (Time, Treatment, Sex × Time, Treatment × Time, Treatment × Sex, Treatment × Sex × Time) (**Figure 1A**).

We next assessed ALT-induced changes in lymphoid populations, including innate lymphoid cells (ILCs), at ZT0 and ZT12 in both sexes (**Figure 1B**). In females, ALT treatment significantly increased ILC2, ILC3, Th1, Th2, and Th17 cells at ZT0 compared with ZT12. In males, ALT exposure similarly increased ILC2 and Th2 cells at ZT0, along with a significant increase in ILC1 cells, whereas ILC3 numbers were significantly higher at ZT12 than at ZT0 following ALT exposure. Baseline Th1 cells in males were higher at ZT0 than ZT12 in PBS-treated controls but were reduced following ALT challenge. In females, ALT exposure induced a marked increase in Th17 cells at ZT0 compared with both PBS-treated controls and ALT-treated mice at ZT12. No significant time-of-day difference in ALT-induced Th17 responses was observed in males. Finally, B cell numbers were significantly increased following ALT challenge at ZT0 in males only, whereas in females ALT exposure resulted in a significant reduction in B cells at ZT12 compared with ZT0 (**Figure 1B**).

Additional interaction analyses revealed significant effects for ILC1, Th1, and B cells across Time, Sex, Treatment, Sex × Time, Treatment × Time, Treatment × Sex, and Treatment × Sex × Time. ILC2 responses were significantly influenced by Time and Treatment, whereas ILC3 responses showed significant effects of Time, Sex, Treatment, and Sex × Time. Th2 cells were significantly affected by Sex and Treatment, while Th17 cells demonstrated significant effects of Sex, Treatment, and Sex × Time (**Figure 1B**).

### Alternaria exposure induces time-of-day-dependent lung inflammation in histopathological analysis

Histopathological assessment of lung sections stained with hematoxylin and eosin (H&E) supported the BAL fluid immune infiltration findings. In female mice, ALT exposure resulted in significantly higher overall lung inflammation scores at ZT12 compared with ZT0, whereas male mice did not exhibit a significant time-of-day difference in inflammation scores (**Figure 2**).

**Figure 2.**
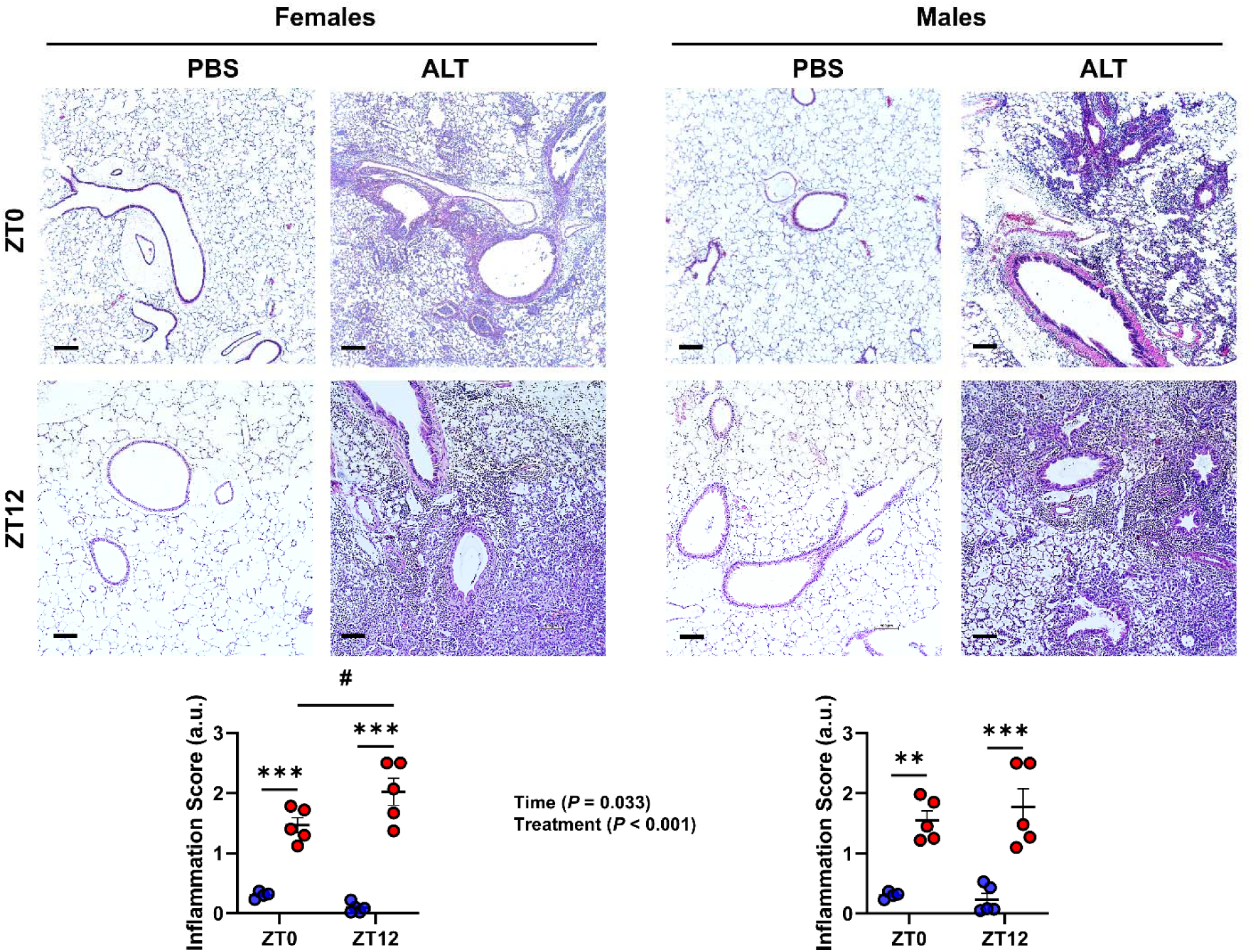
*Alternaria alternata* (ALT) exposure induces time-of-day–dependent lung inflammation as shown by histopathological analysis. Representative Hematoxylin and Eosin (H&E)–stained lung sections illustrating airway inflammation in PBS-treated versus ALT-exposed mice. Data represent mean ± SEM (n = 4–5/group). Statistical analysis was performed using two-way ANOVA followed by Tukey’s multiple comparison test. \*\**P* < 0.01, *** *P* < 0.001 compared with the respective PBS control at ZT0 or ZT12; ^#^ *P* < 0.05, between ALT-exposed groups at ZT0 vs. ZT12. Interactions among Treatment x Sex x Time were analyzed using generalized linear modeling in R (see Supplemental Table S3).

Regional analysis of inflammation further revealed that peribronchial and perivascular inflammation tended to be higher at ZT12 than ZT0 in both females and males, while inflammation within the alveolar region remained comparable between time points (**Figure S1**).

Interaction analyses demonstrated significant effects for overall inflammation scores driven by Time and Treatment. In the alveolar compartment, inflammation scores were significantly influenced by Time, whereas inflammation surrounding the airways and vasculature showed significant effects of Treatment (**Figure 2; Figure S1**).

Collectively, these histopathological findings, together with the BAL cellular analyses, indicate that ALT-induced pulmonary inflammation displays pronounced time-of-day and sex-dependent differences in mice.

### Alternaria-induced airway hyperresponsiveness varies with time of exposure

Based on prior studies demonstrating circadian regulation of allergen-induced airway hyperresponsiveness (AHR) ^28^, we investigated whether the timing of ALT exposure influences AHR severity. Wild-type female and male mice were exposed to ALT for four consecutive days, followed by assessment of airway mechanics using flexiVent in response to a dose-dependent methacholine challenge.

ALT-exposed mice exhibited maximal changes in mean respiratory system resistance (Rrs), elastance (Ers), compliance (Crs), Newtonian resistance (Rn), and tissue damping (G) at the highest methacholine dose (90 mg/mL). When females and males were analyzed together, ALT-exposed mice demonstrated significantly increased Rrs, Ers, Rn, and G at ZT0 compared with ZT12. In contrast, Crs was significantly reduced at ZT0 relative to ZT12, indicating greater airway stiffness during the resting phase (**Figure 3**).

**Figure 3.**
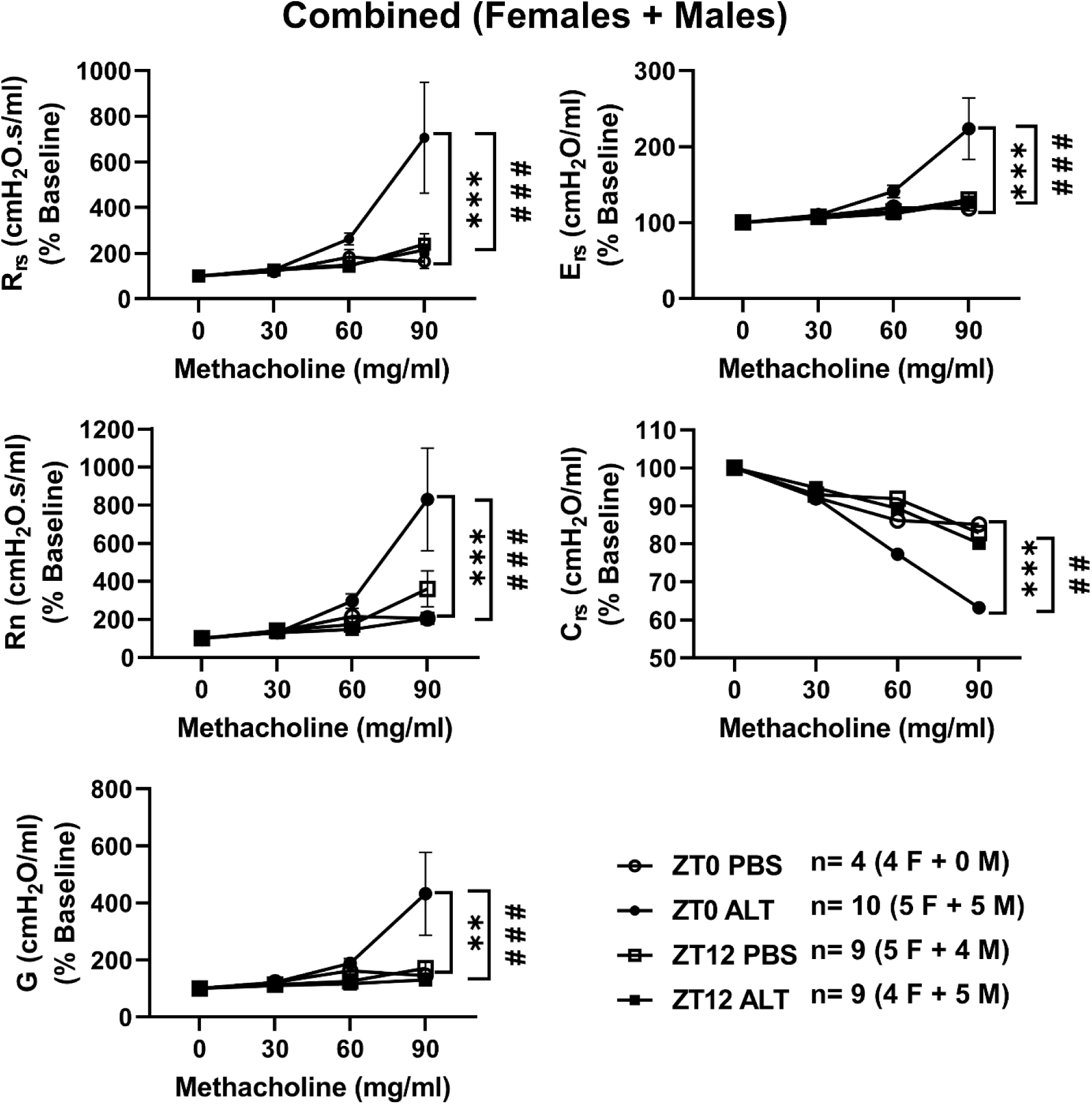
*Alternaria alternata* (ALT) exposure produces time-of-day– and sex-dependent differences in lung function measured by the forced oscillation technique. C57BL/6 wild-type female and male mice were administered ALT extract daily for four consecutive days, and lung function was assessed 24 hours after the final PBS or ALT exposure using the FlexiVent system. Forced oscillation technique parameters were normalized to baseline values at each concentration of aerosolized methacholine in PBS-exposed mice at ZT0 and ZT12 (open circles and open squares) and ALT-exposed mice at ZT0 and ZT12 (closed circles and closed squares). Values were normalized to individual baselines. Data represent mean ± SEM (n=4-10/group; combined analysis of females and males). Statistical analysis was performed using two-way ANOVA followed by Tukey’s multiple comparison test. * * *P* < 0.01, * * * *P* < 0.01, PBS ZT0 vs ALT ZT0 and ^#^ ^#^ *P* < 0.01, ^#^ ^#^ ^#^*P* < 0.001, ALT ZT0 vs ALT ZT12.

Sex-stratified analyses revealed that male mice largely recapitulated these time-of-day-dependent effects, showing significantly greater AHR at ZT0 compared with ZT12 across multiple airway parameters. In contrast, female mice exhibited minimal time-of-day variation in AHR, with the exception of Rn, which was significantly higher at baseline at ZT12 compared with ZT0 (**Figure S2A-B**).

Together, these findings demonstrate that acute ALT exposure induces airway hyperresponsiveness that is modulated by both time of day and sex, with more pronounced circadian variation observed in males than females.

### Alternaria-induced proinflammatory mediators in BAL fluid and lung gene expression display time-of-day- and sex-dependent differences

We have previously shown that ALT exposure induces immune cell infiltration into the lung in a time-of-day– and sex-dependent manner. To determine whether these differences were reflected in inflammatory mediator production, we next quantified cytokines and chemokines in BAL fluid using a custom-designed cytometric bead array.

ALT exposure robustly induced Th2 cytokines in BAL fluid. In females, IL-4, IL-5, and IL-13 levels were significantly increased at both ZT0 and ZT12, with a trend toward higher expression at ZT12. In contrast, males exhibited significantly greater induction of these Th2 cytokines at ZT0 compared with ZT12 (**Figure 4**). Similarly, IL-12p70, IP-10, and IL-6, along with macrophage-associated mediators (TNFα and IL-1β) and Th17 cytokines (IL-17A, IL-17F, and IL-22), were significantly elevated at ZT12 in females. In males, these same mediators, including IP-10, IL-6, TNFα, IL-1β, IL-17A, IL-17F, and IL-22, were significantly higher at ZT0 than at ZT12 (**Figure 4**).

**Figure 4.**
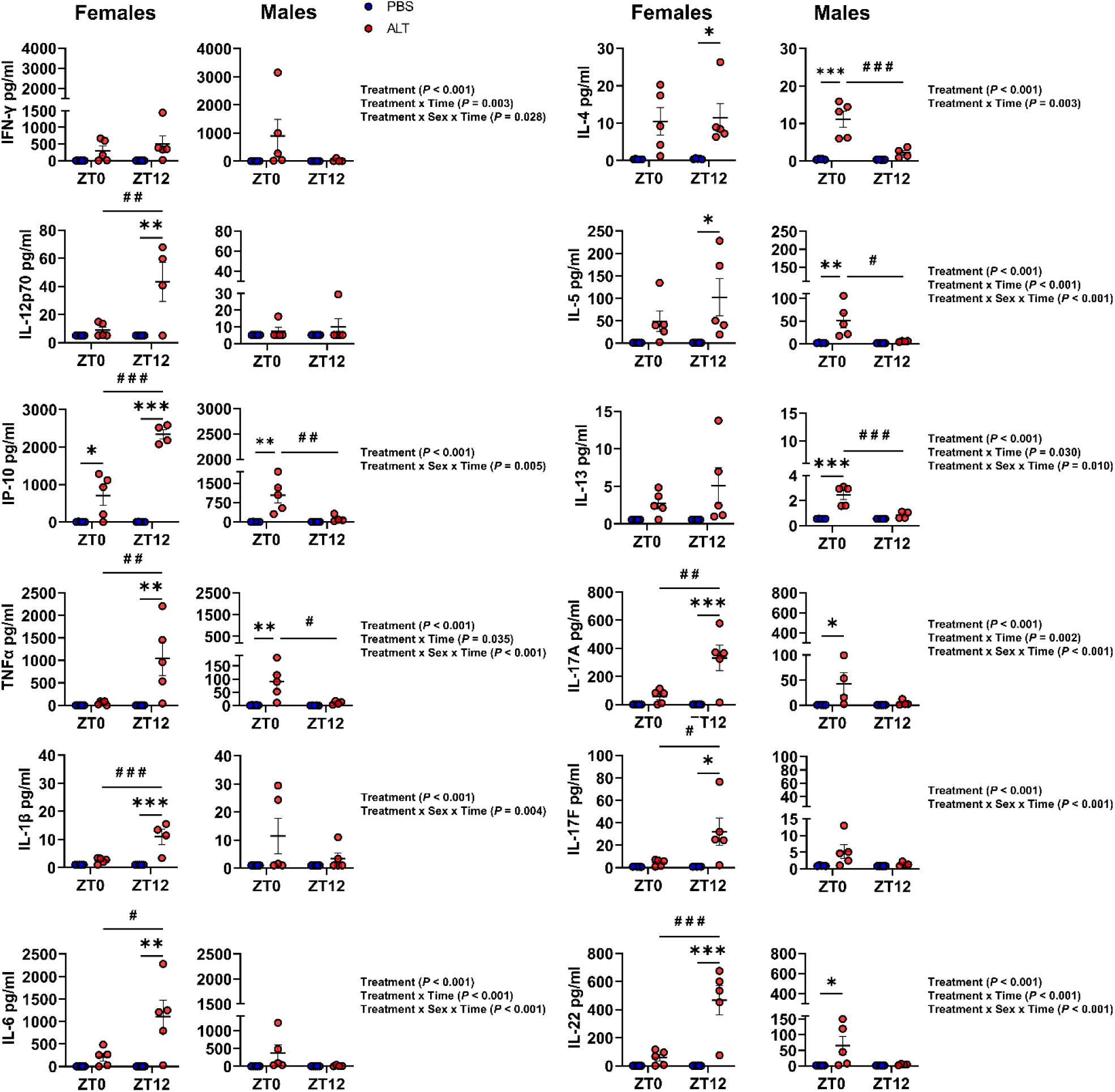
*Alternaria alternata* (ALT) exposure elicits time-of-day and sex-dependent differences in pro-inflammatory mediators in the lung. Wild-type (WT) female and male mice were administered ALT extract daily for four consecutive days. Changes in BALF cytokine/chemokine release were assessed in PBS- and ALT-exposed female and male mice. Pro-inflammatory mediators in BALF were quantified using a cytometric bead array (custom-designed LEGENDplex Mouse Proinflammatory Cytokine/Chemokine panel). Data represent mean ± SEM (n = 4–5/group). Statistical analysis was performed using two-way ANOVA followed by Tukey’s multiple comparison test. \**P* < 0.05, ** *P* < 0.01, *** *P* < 0.001 compared with the respective PBS control at ZT0 or ZT12; ^#^ *P* < 0.05, ^#^ ^#^ *P* < 0.01, ^#^ ^#^ ^#^ *P* < 0.001 between ALT-exposed groups at ZT0 vs. ZT12. Interactions among Treatment x Sex x Time were analyzed using generalized linear modeling in R (see Supplemental Table S3).

Levels of the chemoattractants KC, RANTES, and MCP-1 were significantly increased at ZT12 in females, whereas in males these mediators were elevated at ZT0, demonstrating an opposing circadian pattern between sexes. The epithelial alarmin IL-33 was significantly higher at ZT0 than ZT12 in both females and males following ALT exposure (**Figure S3**). In addition, markers of airway injury and alveolar–capillary barrier disruption, including total BAL protein, uric acid, and lactate dehydrogenase (LDH), were significantly increased at ZT12 compared with ZT0 in females, whereas males showed the inverse pattern, with higher levels at ZT0 (**Figure 5**).

**Figure 5.**
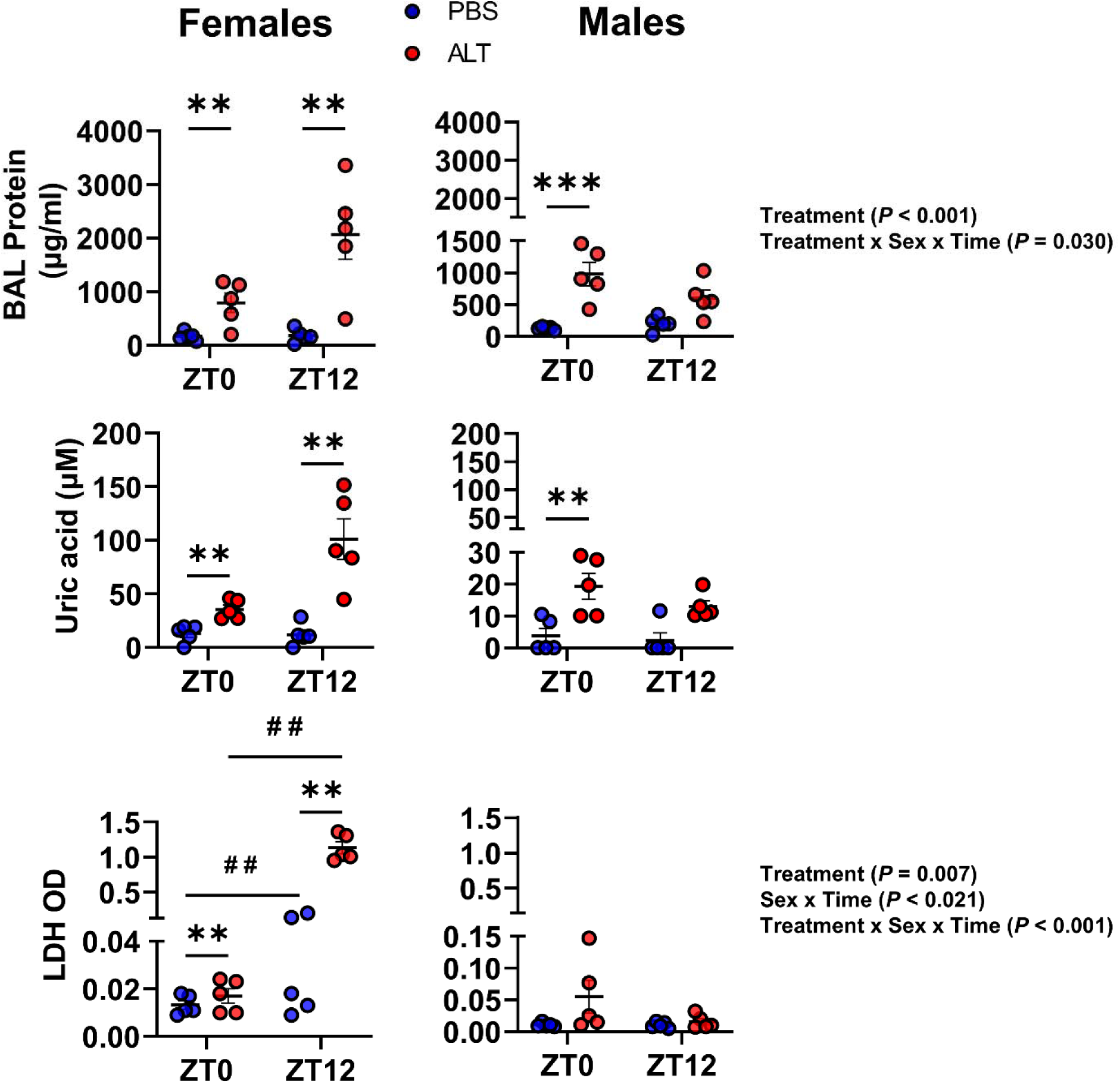
*Alternaria alternata* (ALT) exposure induces time-of-day and sex-dependent differences in barrier integrity and cytotoxicity markers in BAL fluid. Wild-type (WT) female and male mice were administered ALT extract daily for four consecutive days. Total protein in BALF was quantified using a BCA assay, and cytotoxicity markers (uric acid and lactate dehydrogenase) were measured by ELISA in PBS- and ALT-exposed mice. Data represent mean ± SEM (n = 4–5/group). Statistical analysis was performed using two-way ANOVA followed by Tukey’s multiple comparison test. ** *P* < 0.01, *** *P* < 0.001 compared with the respective PBS control at ZT0 or ZT12; ^#^ ^#^ *P* < 0.01 between ALT-exposed groups at ZT0 vs. ZT12. Interactions among Treatment x Sex x Time were analyzed using generalized linear modeling in R (see Supplemental Table S3).

Interaction analyses revealed significant effects across multiple mediators. IFNγ, TNFα, IL-5, IL-6, IL-13, IL-17A, and IL-22 exhibited significant Treatment, Treatment × Time, and Treatment × Sex × Time interactions. IP-10, IL-1β, IL-17F, and total protein showed significant Treatment and Treatment × Sex × Time effects. IL-4 demonstrated significant Treatment and Treatment × Time interactions, while KC exhibited a significant Treatment × Sex × Time interaction. MCP-1 and RANTES were influenced by Treatment, Treatment × Time, and Treatment × Sex × Time. IL-10 showed significant effects of Time, Treatment × Time, and Treatment × Sex × Time, whereas LDH demonstrated significant Treatment, Sex × Time, and Treatment × Sex × Time effects (**Figure 4–5; Figure S3**).

To determine whether these protein-level changes were mirrored at the transcriptional level, we next examined lung mRNA expression of selected proinflammatory cytokines and chemokines. In ALT-exposed females, transcript levels of *Ifn*γ, *Tnf*α, *Il6*, *Il10*, *Il13*, and *Cxcl1* (KC) were significantly higher at ZT12 compared with ZT0 (**Figure 6**). In contrast, ALT-exposed males demonstrated increased *Il4* expression at ZT0, with reduced expression at ZT12. *Il10* expression in males was elevated at ZT0 and remained higher at ZT12, although this difference did not reach statistical significance. *Il13* transcripts were significantly increased in males at both ZT0 and ZT12 compared with respective PBS controls, while expression of other genes remained largely unchanged (**Figure 6**).

**Figure 6.**
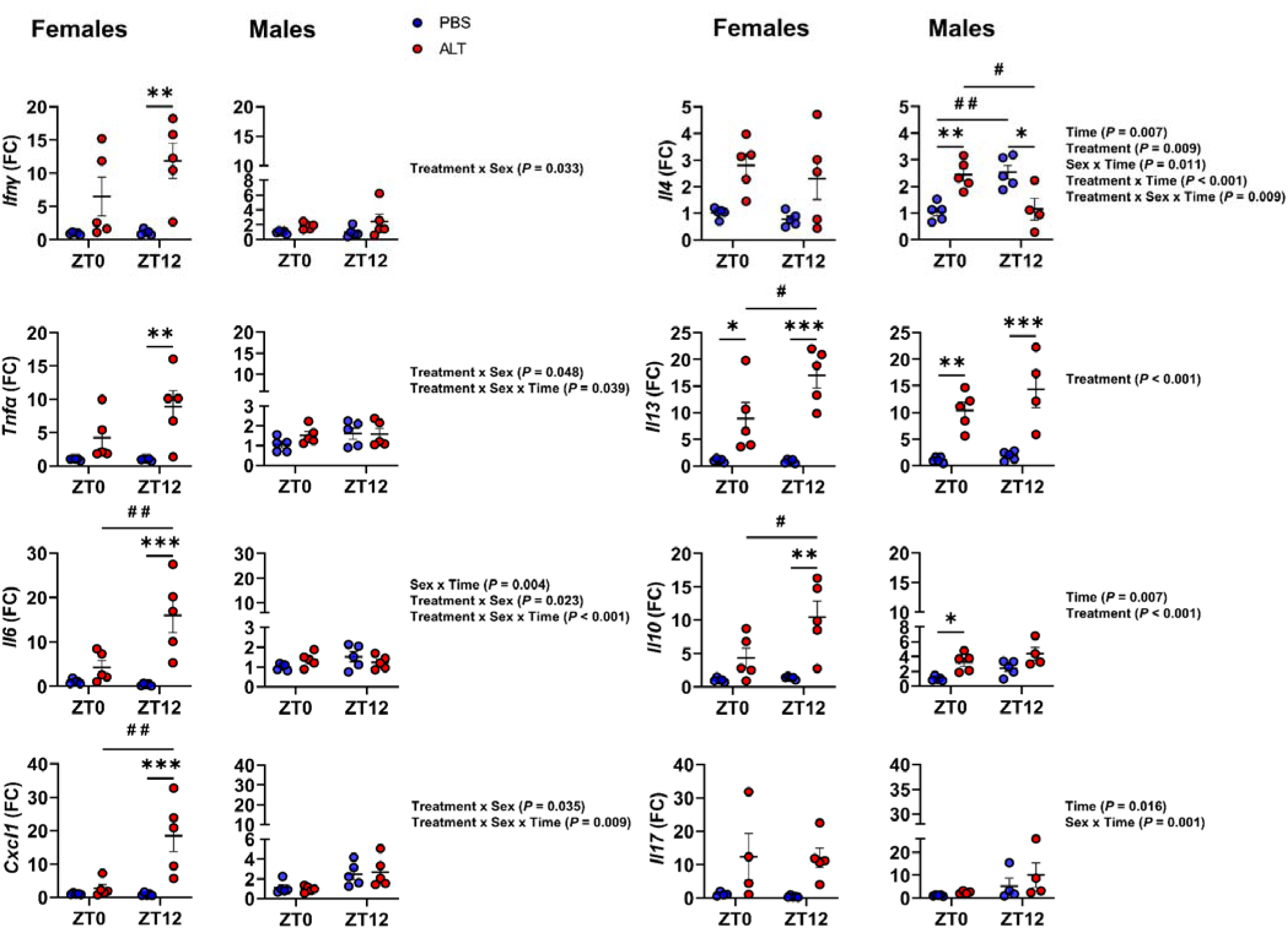
*Alternaria alternata* (ALT) exposure elicits time-of-day and sex-dependent differences in pro-inflammatory gene expression in the lung. Wild-type (WT) female and male mice were administered ALT extract daily for four consecutive days. Changes in transcript levels of pro-inflammatory genes were assessed in PBS- and ALT-exposed female and male mice. Transcript levels of pro-inflammatory genes in lung tissue were measured by quantitative real-time qRT-PCR and normalized to Rplp0 as the housekeeping gene. Data represent mean ± SEM (n = 4–5/group). Statistical analysis was performed using two-way ANOVA followed by Tukey’s multiple comparison test. \**P* < 0.05, ** *P* < 0.01, *** *P* < 0.001 compared with the respective PBS control at ZT0 or ZT12; ^#^ *P* < 0.05, ^#^ ^#^ *P* < 0.01, ^#^ ^#^ ^#^ *P* < 0.001 between ALT-exposed groups at ZT0 vs. ZT12. Interactions among Treatment x Sex x Time were analyzed using generalized linear modeling in R (see Supplemental Table S3).

Further interaction analyses revealed significant effects for *Ifn*γ (Treatment × Sex), *Tnf*α and *Cxcl1* (Treatment × Sex and Treatment × Sex × Time), *Il6* (Sex × Time, Treatment × Sex, and Treatment × Sex × Time), *Il4* (Time, Treatment, Sex × Time, Treatment × Time, and Treatment × Sex × Time), *Il10* (Time and Treatment), *Il13* (Treatment), and *Il17* (Time and Sex × Time) (**Figure 6**).

Collectively, these data demonstrate that ALT exposure elicits pronounced time-of-day– and sex-dependent differences in proinflammatory mediator release in BAL fluid and in inflammatory gene expression within the lung.

### Alternaria exposure differentially regulates circadian clock and epithelial barrier proteins through time-of-day-dependent activation of STAT signaling in the lung

To determine whether the timing of ALT exposure disrupts circadian clock machinery and epithelial barrier integrity in the lung, we performed immunoblot analyses of whole-lung homogenates collected at ZT0 and ZT12.

In ALT-exposed females, protein levels of the core clock components CLOCK and BMAL1 were significantly reduced at ZT0 and remained reduced at ZT12 compared with PBS-treated controls (**Figure 7A**). REV-ERBα protein expression was significantly lower at ZT12 relative to ZT0 following ALT exposure, whereas PER2 showed a modest but nonsignificant reduction at both time points. In males, ALT exposure resulted in significantly reduced CLOCK and BMAL1 expression at ZT12 compared with ZT0, while PER2 levels remained unchanged. In contrast to females, REV-ERBα levels in males were significantly increased at ZT0 but markedly reduced at ZT12 following ALT exposure (**Figure 7A**). These sex-specific changes were corroborated at the transcriptional level, as *Rev-erb*α mRNA expression exhibited a similar time-of-day–dependent pattern (**Figure S4A**).

**Figure 7.**
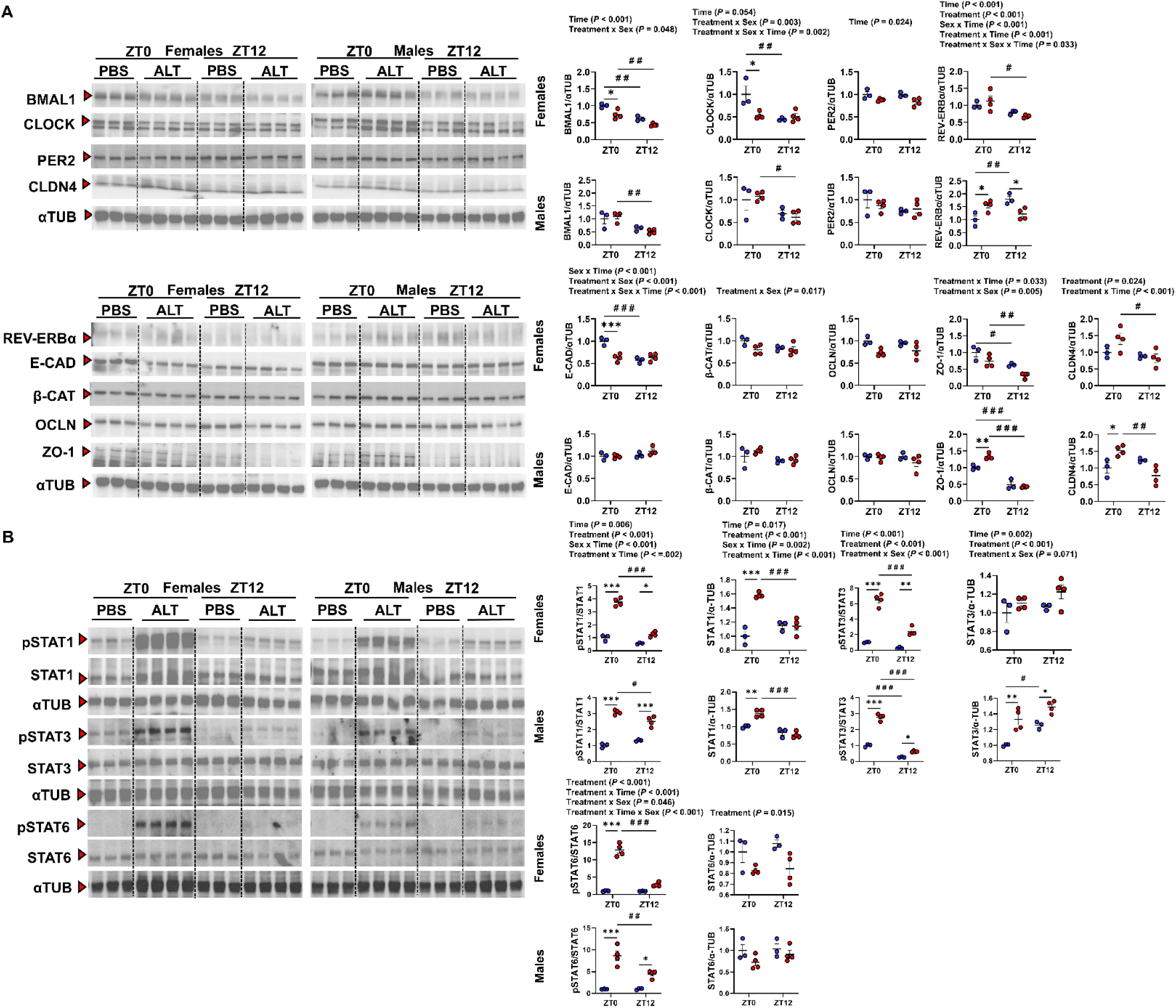
*Alternaria alternata* (ALT) exposure alters protein abundance of circadian clock and epithelial barrier components, potentially through STAT activation. (A–B) ALT exposure induces time-of-day–dependent changes in the protein expression of circadian clock regulators, epithelial barrier markers, and STAT signaling molecules in female and male mice. Representative Western blot images show alterations in core circadian clock proteins, epithelial barrier proteins, and STAT family members in PBS- and ALT-treated groups at ZT0 and ZT12. α-Tubulin was used as the loading control for densitometric analysis. Data represent mean ± SEM (n = 3–4/group). Statistical analysis was performed using two-way ANOVA followed by Tukey’s multiple comparison test. \**P* < 0.05, \*\**P* < 0.01, \*\*\**P* < 0.001 compared with the respective PBS control at ZT0 or ZT12; ^#^*P* < 0.05, ^#^ ^#^*P* < 0.01, ^#^ ^#^ ^#^*P* < 0.001 between ALT-exposed groups at ZT0 vs. ZT12. Interactions among Treatment x Sex x Time were analyzed using generalized linear modeling in R (see Supplemental Table S3).

Interaction analyses revealed significant effects for BMAL1 protein expression (Time and Treatment × Sex), REV-ERBα (Time, Treatment, Sex × Time, Treatment × Time, and Treatment × Sex × Time), CLOCK (Time, Treatment × Time, and Treatment × Sex × Time), and PER2 (Time). Significant interactions were also observed for *Rev-erb*α transcript levels (Time, Treatment, Sex × Time, and Treatment × Time) (**Figure 7A**).

We next assessed whether ALT exposure altered pulmonary epithelial barrier integrity. In females, ALT exposure led to a time-dependent reduction in E-CADHERIN and ZO-1 expression, with significantly higher ZO-1 levels observed at ZT0 than ZT12 compared with ALT-treated group. ALT exposure reduced ZO-1 levels at both time points, whereas E-CADHERIN was reduced predominantly at ZT0. In addition, β-CATENIN and OCCLUDIN expression were reduced by ALT at both ZT0 and ZT12, whereas PBS-treated controls showed no time-of-day differences. CLAUDIN-4 expression was modestly increased by ALT at ZT0 but significantly reduced at ZT12 (**Figure 7A**).

In ALT-exposed males, no significant time-of-day differences were observed in E-CADHERIN, β-CATENIN, or OCCLUDIN expression. However, ZO-1 expression in PBS-treated males was significantly higher at ZT0 than ZT12, and ALT exposure further increased ZO-1 levels at ZT0 while remaining unchanged at ZT12 compared with controls. Similar to females, CLAUDIN-4 expression in males increased at ZT0 but decreased at ZT12 following ALT exposure. Interaction analyses revealed significant effects for E-CADHERIN (Sex × Time, Treatment × Sex, Treatment × Sex × Time), β-CATENIN (Treatment × Sex), ZO-1 (Treatment × Time and Treatment × Sex), and CLAUDIN-4 (Treatment and Treatment × Time) (**Figure 7A**).

Consistent with these protein-level findings, analysis of epithelial barrier gene expression demonstrated that *Cdh1* and *Ctnnb1* transcripts showed an increasing trend following ALT exposure at ZT12 in both sexes. In contrast, *Ocln* transcript levels were significantly higher at ZT0 than ZT12 in females, whereas in males *Ocln* expression was higher at ZT12 (**Figure S4B**).

Given prior evidence linking ALT exposure to STAT pathway activation ^29^, we next examined whether time-dependent immune and epithelial responses were associated with differential STAT signaling. ALT exposure markedly increased phosphorylation of STAT1, STAT3, and STAT6 at ZT0 compared with ZT12 in both females and males (**Figure 7B**).

Among total STAT proteins, STAT1 expression was significantly higher at ZT0 than ZT12 following ALT exposure in both sexes. In males, total STAT3 expression was higher at ZT12 than ZT0 in both PBS- and ALT-treated mice, whereas females exhibited no time-of-day differences in total STAT3. Total STAT6 levels remained unchanged in both sexes at ZT0 and ZT12 (**Figure 7B**). Consistent with these results, no significant changes were detected in *Stat3* or *Stat6* transcript levels across time points or treatment groups (**Figure S4C**).

Interaction analyses revealed significant effects for phosphorylation of STAT1 (Time, Treatment, Sex × Time, Treatment × Sex, Treatment × Time), STAT3 (Time, Treatment, Treatment × Sex), and STAT6 (Treatment, Treatment × Time, Treatment × Sex, Treatment × Sex × Time) (**Figure 7B**).

Since STAT signaling can crosstalk with MAPK pathways, we also examined ERK activation. Following ALT exposure, phosphorylation of ERK1/2 at ZT0 was reduced in females but significantly increased in males, whereas no changes were observed at ZT12 in either sex (**Figure S5**). Total ERK1/2 protein levels were significantly higher at ZT12 than ZT0 in ALT-exposed females but remained unchanged in males at both time points. Interaction analyses showed significant effects for phosphorylated ERK1/2 (Treatment, Sex × Time, Treatment × Time, Treatment × Sex, Treatment × Sex × Time) and total ERK1/2 (Sex × Time, Treatment × Time, Treatment × Sex, Treatment × Sex × Time) (**Figure S5**).

Together, these findings indicate that ALT exposure disrupts circadian clock regulation and epithelial barrier integrity in a time-of-day– and sex-dependent manner, potentially mediated through differential activation of STAT and ERK signaling pathways in the lung.

## Discussion

Circadian regulation of leukocyte trafficking is well established, with immune cell homing governed by rhythmic oscillations in chemokines and adhesion molecules ^30, 31^. Accordingly, immune response to environmental allergens varies with time of day, as shown by our group and others ^27, 28, 32^. Chronic inflammatory lung diseases such as asthma are also under circadian control, with disease severity, exacerbations, and mortality peaking during late night or early morning hours ^6, 7^. Exposure to *Alternaria alternata* (ALT), a ubiquitous fungal allergen, is strongly associated with severe asthma symptoms, hospitalization, and fatal attacks, particularly among children and young adults, and is linked to persistent airflow obstruction and disease severity ^8, 9, 11, 12^.

Building upon these observations, we investigated whether circadian timing directly modulates inflammatory responses to fungal allergen exposure and whether these responses differ from those reported for house dust mite (HDM) ^27, 28, 32^. Our findings demonstrate that ALT induces robust allergic airway inflammation, airway hyperresponsiveness, and epithelial barrier disruption in a time-of-day- and sex-dependent manner. The marked influx of immune cells into the bronchoalveolar lavage fluid at both time points confirms the strong immunogenicity of ALT, consistent with its clinical association with severe asthma exacerbations ^8, 9^. Notably, females displayed a distinct temporal immune profile, characterized by increased eosinophils (GR1^+^ and inflammatory subsets), neutrophils, interstitial macrophages and dendritic cells) at ZT0 (rest phase), whereas males exhibited greater inflammatory infiltration at ZT12 (active phase), with dendritic cells showing less temporal variation.

This sex-specific temporal pattern contrasts with our prior HDM findings, where male mice showed increased granulocyte infiltration at ZT12, while females exhibit heightened but differing responses depending on cell type ^27^. Although neutrophil and dendritic cell responses and reductions in alveolar macrophages were largely similar between HDM and ALT exposure, the divergent eosinophilic and macrophage patterns suggest that ALT reshapes lung immune composition in a temporally and sexually dimorphic manner. These differences may reflect hormone-dependent modulation of circadian immune responses, as previously suggested ^33–37^

Given the established role of epithelial-derived alarmins in ALT-induced inflammation, we next examined innate lymphoid and T helper cell responses. Consistent with prior studies ^24, 35, 37^, ALT exposure increased IL-33 release and promoted accumulation of ILC2 and Th2 cells in both sexes. Females showed concordant increases in ILC2s, Th2 cells and eosinophils at ZT0, whereas males displayed a temporal dissociation, with lymphoid cell expansion at ZT0 but peak eosinophilic infiltration at ZT12. In contrast, ILC1, ILC3, Th1, and Th17 populations exhibited opposing sex- and time-dependent patterns, suggesting temporal segregation of immune pathways that may optimize host defense. To our knowledge, this is the first study to define time-of-day-dependent lymphoid responses to ALT exposure in a murine asthma model.

Cytokine and chemokine production by epithelial and immune cells is also regulated by the circadian clock ^38, 39^. ALT exposure induced robust Th2 proinflammatory, and Th17 cytokine responses that varied with exposure time and sex. Females exhibited higher levels of IL-4, IL-5, IL-13, IL-12p70, IP-10, IL-6, TNFα, IL-1β, IL-17A, IL-17F, and IL-22 at ZT12, whereas males showed maximal responses at ZT0. Similar temporal patterns were observed for chemokines KC, RANTES, and MCP-1. These findings differ from prior HDM models, where peak inflammation consistently occurred at ZT12 ^27, 32^, highlighting allergen-specific differences likely influenced by exposure duration and innate sensing mechanisms. Transcript-level analyses largely mirrored protein expression in females, while males showed limited circadian variation, apart from increased *Il4* expression at ZT0. Markers of tissue damage, including lactate dehydrogenase and uric acid, followed these same circadian-dependent inflammatory patterns ^24, 35^.

Functionally, ALT-exposed mice exhibited greater airway hyperresponsiveness during the resting phase consistent with clinical observations of circadian variation in asthma severity ^6, 7, 28^. Increased airway resistance, elastance, tissue damping, and reduced compliance at ZT0 indicated heightened airway stiffness. Sex-stratified analyses revealed that males largely accounted for this circadian variation, whereas females showed minimal diurnal differences in airway mechanics despite pronounced inflammatory responses. These findings suggest sex-specific uncoupling between airway inflammation and functional obstruction.

Epithelial barrier integrity is essential for protecting the lung from inhaled allergens and pathogens ^40^. Consistent with our prior studies showing allergen-mediated disruption of tight junction proteins ^41, 42^, ALT exposure reduced E-cadherin, ZO-1, and occludin expression in females, particularly at ZT0, while males showed minimal changes. Claudin-4 expression exhibited circadian-dependent alterations in both sexes. Collectively, these findings indicate compromised epithelial barrier function, a hallmark of allergic asthma that facilitates allergen penetration and amplifies inflammation ^40^. Importantly, circadian clocks regulate epithelial junctional complexes ^43–45^ and circadian disruption has been linked to asthma severity in both experimental models and human tissues ^27, 28, 45–47^. Our observation that ALT alters expression of core clock proteins CLOCK, BMAL1, and REV-ERBα suggests that fungal allergens may directly perturb lung circadian rhythms, contributing to barrier dysfunction and immune dysregulation.

Mechanistically, STAT signaling emerged as a key pathway linking circadian regulation with immune and epithelial responses. Enhanced phosphorylation of STAT1, STAT3, and STAT6 at ZT0 coincided with peak inflammatory responses, particularly in males, consistent with established roles for these pathways in Th2 and Th17 polarization ^18, 29, 48, 49^. Given evidence that circadian clock components regulate STAT signaling ^50^, these findings support a bidirectional interaction between molecular clocks and inflammatory pathways. To our knowledge, this is the first report demonstrating time-of-day–dependent STAT activation following fungal allergen exposure in the lung.

This study has limitations. We did not assess sex hormone levels or evaluate epithelial versus immune cell–specific contributions to barrier dysfunction and STAT activation using complementary *in vitro* models. Additionally, our findings are based on an acute ALT exposure model; responses may differ with chronic or repeated exposure. Future studies will examine these variables to define circadian mechanisms driving asthma exacerbations.

## Conclusions

In summary, our study demonstrates that ALT-induced allergic airway inflammation is regulated by circadian timing and sex through interconnected epithelial, immune, and clock-controlled signaling pathways. These findings highlight the importance of considering time-of-day and biological sex in asthma pathobiology and support the potential for chronobiology-informed therapeutic strategies to improve asthma management.

## CRediT Authorship Contribution Statement

I.K.S. and S.K.D. conceptualized and designed the study. S.K.D. and I.K.S. were primarily responsible for experimental design, data analysis, critical interpretation of the data, figure preparation, and manuscript drafting. J.M. and S.K.D. performed immunoblot analyses and interpreted the results. S.K.D., C.P., and I.K.S. conducted all mouse experiments. S.H. performed biostatistical interaction analyses. S.K.D., J.M., C.P., S.H., M.C., and I.K.S. reviewed the manuscript and approved the final version.

## Funding

This work was supported in part by the National Institutes of Health (NIH) grant R01HL142543 (to I.K.S.) and by startup funds from the University of Kansas Medical Center, School of Medicine, Department of Internal Medicine (to I.K.S.). The content is solely the responsibility of the authors and does not necessarily represent the official views of the National Institutes of Health.

## Acknowledgements

SMART-Servier Medical Art (https://smart.servier.com/) was utilized in creating the graphical abstract.

## Declaration of Competing Interest

The authors declare no conflicts of interest or personal relationships that could have influenced the work reported in this manuscript.

## Appendix A. Supplementary Materials

Online Supplementary Methods

Supplementary Figures S1-S5

Supplementary Tables S1-S3

## Ethics approval and consent to participate

All animal experiments were conducted in accordance with institutional guidelines and were approved by the University of Kansas Medical Center Institutional Animal Care and Use Committee (IACUC) (protocol number ACUP #2020-2575-1).

## Data and Material Availability

The data supporting the findings of this study are available from the corresponding author upon reasonable request.

## Notes

### Competing Interest Statement

The authors have declared no competing interest.

## References

1. Mohawk JA, Green CB, Takahashi JS. Central and peripheral circadian clocks in mammals. Annu Rev Neurosci 2012; 35: 445–462. 10.1146/annurev-neuro-060909-153128

2. Giri A, Wang Q, Rahman I, Sundar IK. Circadian molecular clock disruption in chronic pulmonary diseases. Trends Mol Med 2022; 28(6): 513–527. 10.1016/j.molmed.2022.04.002

3. Sundar IK, Yao H, Sellix MT, Rahman I. Circadian molecular clock in lung pathophysiology. Am J Physiol Lung Cell Mol Physiol 2015; 309(10): L1056–1075. 10.1152/ajplung.00152.2015

4. Cribb L, Sha R, Yiallourou S, Grima NA, Cavuoto M, Baril A-A et al. Sleep regularity and mortality: a prospective analysis in the UK Biobank. eLife 2023; 12: RP88359. 10.7554/eLife.88359

5. Das MK, De Ryck E, Jorgensen IL, Zienolddiny-Narui S, Erdem JS. Circadian rhythm disruption in cardiovascular disease: a systematic review and meta-analysis of mechanistic evidence from animal models. BMC Med 2026; 24(1): 73 10.1186/s12916-025-04572-3

6. Scheer F, Hilton MF, Evoniuk HL, Shiels SA, Malhotra A, Sugarbaker R et al. The endogenous circadian system worsens asthma at night independent of sleep and other daily behavioral or environmental cycles. Proc Natl Acad Sci U S A 2021; 118(37). 10.1073/pnas.2018486118

7. Troyanov S, Ghezzo H, Cartier A, Malo JL. Comparison of circadian variations using FEV1 and peak expiratory flow rates among normal and asthmatic subjects. Thorax 1994; 49(8): 775–780. 10.1136/thx.49.8.775

8. Medrek SK, Kao CC, Yang DH, Hanania NA, Parulekar AD. Fungal Sensitization Is Associated with Increased Risk of Life-Threatening Asthma. J Allergy Clin Immunol Pract 2017; 5(4): 1025–1031.e1022. 10.1016/j.jaip.2016.11.015

9. Masaki K, Fukunaga K, Matsusaka M, Kabata H, Tanosaki T, Mochimaru T et al. Characteristics of severe asthma with fungal sensitization. Ann Allergy Asthma Immunol 2017; 119(3): 253–257. 10.1016/j.anai.2017.07.008

10. Vailes LD, Perzanowski MS, Wheatley LM, Platts-Mills TA, Chapman MD. IgE and IgG antibody responses to recombinant Alt a 1 as a marker of sensitization to Alternaria in asthma and atopic dermatitis. Clin Exp Allergy 2001; 31(12): 1891–1895. 10.1046/j.1365-2222.2001.00745.x

11. Hadebe S, Brombacher F. Environment and Host-Genetic Determinants in Early Development of Allergic Asthma: Contribution of Fungi. Front Immunol 2019; 10: 2696.10.3389/fimmu.2019.02696

12. O’Hollaren MT, Yunginger JW, Offord KP, Somers MJ, O’Connell EJ, Ballard DJ et al. Exposure to an aeroallergen as a possible precipitating factor in respiratory arrest in young patients with asthma. N Engl J Med 1991; 324(6): 359–363. 10.1056/nejm199102073240602

13. Leino MS, Loxham M, Blume C, Swindle EJ, Jayasekera NP, Dennison PW et al. Barrier disrupting effects of alternaria alternata extract on bronchial epithelium from asthmatic donors. PLoS One 2013; 8(8): e71278. 10.1371/journal.pone.0071278

14. Toki S, Goleniewska K, Zhang J, Zhou W, Newcomb DC, Zhou B et al. TSLP and IL-33 reciprocally promote each other’s lung protein expression and ILC2 receptor expression to enhance innate type-2 airway inflammation. Allergy 2020; 75(7): 1606–1617. 10.1111/all.14196

15. Kouzaki H, Iijima K, Kobayashi T, O’Grady SM, Kita H. The danger signal, extracellular ATP, is a sensor for an airborne allergen and triggers IL-33 release and innate Th2-type responses. J Immunol 2011; 186(7): 4375–4387. 10.4049/jimmunol.1003020

16. Grinn-Gofroń A, Bosiacka B. Effects of meteorological factors on the composition of selected fungal spores in the air. Aerobiologia (Bologna) 2015; 31(1): 63–72. 10.1007/s10453-014-9347-1

17. Kuperman D, Schofield B, Wills-Karp M, Grusby MJ. Signal transducer and activator of transcription factor 6 (Stat6)-deficient mice are protected from antigen-induced airway hyperresponsiveness and mucus production. J Exp Med 1998; 187(6): 939–948. 10.1084/jem.187.6.939

18. Simeone-Penney MC, Severgnini M, Tu P, Homer RJ, Mariani TJ, Cohn L et al. Airway epithelial STAT3 is required for allergic inflammation in a murine model of asthma. J Immunol 2007; 178(10): 6191–6199. 10.4049/jimmunol.178.10.6191

19. Shimoda K, van Deursen J, Sangster MY, Sarawar SR, Carson RT, Tripp RA et al. Lack of IL-4-induced Th2 response and IgE class switching in mice with disrupted Stat6 gene. Nature 1996; 380(6575): 630–633. 10.1038/380630a0

20. Hoshino A, Tsuji T, Matsuzaki J, Jinushi T, Ashino S, Teramura T et al. STAT6-mediated signaling in Th2-dependent allergic asthma: critical role for the development of eosinophilia, airway hyper-responsiveness and mucus hypersecretion, distinct from its role in Th2 differentiation. Int Immunol 2004; 16(10): 1497–1505. 10.1093/intimm/dxh151

21. Xue Y, Zhou Y, Bao W, Fu Q, Hao H, Han L et al. STAT3 and IL-6 Contribute to Corticosteroid Resistance in an OVA and Ozone-induced Asthma Model with Neutrophil Infiltration. Front Mol Biosci 2021; 8: 717962. 10.3389/fmolb.2021.717962

22. Doherty TA, Khorram N, Sugimoto K, Sheppard D, Rosenthal P, Cho JY et al. Alternaria induces STAT6-dependent acute airway eosinophilia and epithelial FIZZ1 expression that promotes airway fibrosis and epithelial thickness. J Immunol 2012; 188(6): 2622–2629. 10.4049/jimmunol.1101632

23. Halwani R, Sultana A, Vazquez-Tello A, Jamhawi A, Al-Masri AA, Al-Muhsen S. Th-17 regulatory cytokines IL-21, IL-23, and IL-6 enhance neutrophil production of IL-17 cytokines during asthma. J Asthma 2017; 54(9): 893–904. 10.1080/02770903.2017.1283696

24. Snelgrove RJ, Gregory LG, Peiró T, Akthar S, Campbell GA, Walker SA et al. Alternaria-derived serine protease activity drives IL-33-mediated asthma exacerbations. J Allergy Clin Immunol 2014; 134(3): 583–592.e586. 10.1016/j.jaci.2014.02.002

25. Hartney JM, Robichaud A. Assessment of airway hyperresponsiveness in mouse models of allergic lung disease using detailed measurements of respiratory mechanics. Methods Mol Biol 2013; 1032: 205–217. 10.1007/978-1-62703-496-8_16

26. Shalaby KH, Gold LG, Schuessler TF, Martin JG, Robichaud A. Combined forced oscillation and forced expiration measurements in mice for the assessment of airway hyperresponsiveness. Respir Res 2010; 11(1): 82 10.1186/1465-9921-11-82

27. Srinivasan A, Giri A, Duraisamy SK, Alsup A, Castro M, Sundar IK. Acute HDM exposure shows time-of-day and sex-based differences in the severity of lung inflammation and circadian clock disruption. J Allergy Clin Immunol Glob 2023; 2(4):100155. 10.1016/j.jacig.2023.100155

28. Durrington HJ, Krakowiak K, Meijer P, Begley N, Maidstone R, Goosey L et al. Circadian asthma airway responses are gated by REV-ERBα. Eur Respir J 2020; 56(6). 10.1183/13993003.02407-2019

29. Valladao AC, Frevert CW, Koch LK, Campbell DJ, Ziegler SF. STAT6 Regulates the Development of Eosinophilic versus Neutrophilic Asthma in Response to Alternaria alternata. J Immunol 2016; 197(12): 4541–4551. 10.4049/jimmunol.1600007

30. Scheiermann C, Kunisaki Y, Frenette PS. Circadian control of the immune system. Nat Rev Immunol 2013; 13(3): 190–198. 10.1038/nri3386

31. Pick R, He W, Chen CS, Scheiermann C. Time-of-Day-Dependent Trafficking and Function of Leukocyte Subsets. Trends Immunol 2019; 40(6): 524–537. 10.1016/j.it.2019.03.010

32. Srinivasan A, Giri A, Duraisamy SK, Alsup A, Castro M, Sundar IK. Chronic HDM exposure shows time-of-day and sex-based differences in inflammatory response associated with lung circadian clock disruption. iScience 2023; 26(9): 107580. 10.1016/j.isci.2023.107580

33. Cephus JY, Stier MT, Fuseini H, Yung JA, Toki S, Bloodworth MH et al. Testosterone Attenuates Group 2 Innate Lymphoid Cell-Mediated Airway Inflammation. Cell Rep 2017; 21(9): 2487–2499. 10.1016/j.celrep.2017.10.110

34. Kozieł MJ, Habrowska-Górczyńska DE, Urbanek KA, Domińska K, Piastowska-Ciesielska AW, Kowalska K. Estrogen receptor α mediates alternariol-induced apoptosis and modulation of the invasiveness of ovarian cancer cells. Toxicology Letters 2023; 386: 9–19.10.1016/j.toxlet.2023.09.001

35. Cephus JY, Gandhi VD, Shah R, Brooke Davis J, Fuseini H, Yung JA et al. Estrogen receptor-α signaling increases allergen-induced IL-33 release and airway inflammation. Allergy 2021; 76(1): 255–268. 10.1111/all.14491

36. Stölting H, Raftery AL, Walker SA, Rutherford EN, Gore ML, Huang Y, et al. Epidermal growth factor receptor controls sex differences in lung type 2 responses to inhaled allergen. Sci Immunol 2026; 11(115): eadk1673. 10.1126/sciimmunol.adk1673

37. Wang C, Xu ZB, Peng YQ, Zhang HY, Yu QN, Guo YB et al. Sex differences in group 2 innate lymphoid cell-dominant allergic airway inflammation. Mol Immunol 2020; 128: 89–97. 10.1016/j.molimm.2020.09.019

38. Gibbs JE, Blaikley J, Beesley S, Matthews L, Simpson KD, Boyce SH et al. The nuclear receptor REV-ERBα mediates circadian regulation of innate immunity through selective regulation of inflammatory cytokines. Proc Natl Acad Sci U S A 2012; 109(2): 582–587. 10.1073/pnas.1106750109

39. Petrovsky N, McNair P, Harrison LC. Diurnal rhythms of pro-inflammatory cytokines: regulation by plasma cortisol and therapeutic implications. Cytokine 1998; 10(4): 307–312. 10.1006/cyto.1997.0289

40. Georas SN, Rezaee F. Epithelial barrier function: at the front line of asthma immunology and allergic airway inflammation. J Allergy Clin Immunol 2014; 134(3): 509–520. 10.1016/j.jaci.2014.05.049

41. Duraisamy SK, Srinivasan A, Sundar IK. House dust mite and Th2 cytokine-mediated epithelial barrier dysfunction attenuation by KL001 in 16-HBE cells. Tissue Barriers 2024; 12(1): 2203841. 10.1080/21688370.2023.2203841

42. Duraisamy SK, Sundar IK. REV-ERBα agonist SR10067 attenuates Th2 cytokine-mediated barrier dysfunction in human bronchial epithelial cells. Clin Sci (Lond) 2024; 138(19): 1209–1226. 10.1042/cs20240064

43. Summa KC, Voigt RM, Forsyth CB, Shaikh M, Cavanaugh K, Tang Y et al. Disruption of the Circadian Clock in Mice Increases Intestinal Permeability and Promotes Alcohol-Induced Hepatic Pathology and Inflammation. PLoS One 2013; 8(6): e67102. 10.1371/journal.pone.0067102

44. Kyoko OO, Kono H, Ishimaru K, Miyake K, Kubota T, Ogawa H et al. Expressions of tight junction proteins Occludin and Claudin-1 are under the circadian control in the mouse large intestine: implications in intestinal permeability and susceptibility to colitis. PLoS One 2014; 9(5): e98016. 10.1371/journal.pone.0098016

45. Nakao A. Clockwork allergy: How the circadian clock underpins allergic reactions. J Allergy Clin Immunol 2018; 142(4): 1021–1031. 10.1016/j.jaci.2018.08.007

46. Ehlers A, Xie W, Agapov E, Brown S, Steinberg D, Tidwell R et al. BMAL1 links the circadian clock to viral airway pathology and asthma phenotypes. Mucosal Immunol 2018; 11(1): 97–111. 10.1038/mi.2017.24

47. Powell WT, Clark LV, Rich LM, Vanderwall ER, Gates C, White MP et al. Altered circadian gene expression in primary human airway epithelial cells in asthma. ERJ Open Res 2024; 10(4). 10.1183/23120541.00005-2024

48. Sampath D, Castro M, Look DC, Holtzman MJ. Constitutive activation of an epithelial signal transducer and activator of transcription (STAT) pathway in asthma. J Clin Invest 1999; 103(9): 1353–1361. 10.1172/jci6130

49. Mullings RE, Wilson SJ, Puddicombe SM, Lordan JL, Bucchieri F, Djukanović R et al. Signal transducer and activator of transcription 6 (STAT-6) expression and function in asthmatic bronchial epithelium. Journal of Allergy and Clinical Immunology 2001; 108(5): 832–838. 10.1067/mai.2001.119554

50. Wang H, Fu Y. NR1D1 suppressed the growth of ovarian cancer by abrogating the JAK/STAT3 signaling pathway. BMC Cancer 2021; 21(1): 871. 10.1186/s12885-021-08597-8

