## Supplemental Methods for "Circadian and Sex-Specific Responses to *Alternaria alternata* in Allergic Airway Inflammation"

**Short Running Title: Circadian Regulation of Fungal Allergen–Induced Asthma**

**Inflammatory Mediators in BAL fluid**

BAL fluid aliquots stored at −80°C were thawed on ice and analyzed for inflammatory mediators using a custom-designed LEGENDplex™ Mouse Proinflammatory Cytokine/Chemokine Panel (BioLegend), following the manufacturer’s instructions. Briefly, 100 µL of BAL fluid was incubated with capture bead populations coated with analyte‑specific antibodies in a V‑bottom plate at 37°C. After incubation, beads were washed and processed through sequential addition of streptavidin‑PE detection reagent, followed by additional wash steps. Samples were acquired on an Attune NxT flow cytometer (Thermo Fisher Scientific), and cytokine/chemokine concentrations were quantified using the LEGENDplex Data Analysis Suite (BioLegend). Cytokine and chemokine concentrations were calculated and expressed as pg/ml, as described previously ^1,2^. The lower limit of detection (LLOD) for each analyte in this assay were as follows: IFNγ (1.38 pg/ml), TNFα (2.31 pg/ml), IL-6 (0.16 pg/ml), IL-4 (0.41 pg/ml), IL-10 (3.341 pg/ml), IL-17A (0.54 pg/ml), IL-17F (1.09 pg/ml), IL-22 (1.96 pg/ml), IL-13 (0.82 pg/ml), KC (0.97 pg/ml), MCP-1 (0.52 pg/ml), IL-12p70 (0.56 pg/ml), RANTES (6.58 pg/ml), IL-1β (0.58 pg/ml), IP-10 (2.78 pg/ml), and GM-CSF (0.33 pg/ml). All values at or close to the LLOD were used as it is for data analysis and comparison of groups although they may not reflect the absolute concentrations.

**Measurement of Total Protein, IL-33, Uric Acid, and LDH in BAL Fluid**

Total protein levels in BAL fluid, used as an indicator of alveolar barrier dysfunction, were quantified using the Pierce™ Micro BCA Protein Assay Kit (Thermo Fisher Scientific, USA) following the manufacturer’s instructions. Alarmin cytokine IL-33 release in BAL fluid measured using ELISA kit (Invitrogen, Thermo Fisher Scientific, USA). To assess Alternaria‑induced cytotoxicity, uric acid and lactate dehydrogenase (LDH) levels in BAL fluid were measured using Amplex Red Uric Acid/Uricase Assay Kit (Molecular Probes, Life Technologies, Thermo Fisher Scientific, USA) and CyQUANT™ LDH Cytotoxicity Assay Kit (Invitrogen, Thermo Fisher Scientific, USA). All assays were performed according to the manufacturers’ protocols, and absorbance or fluorescence values were recorded using standard microplate‑based detection methods.

**Protein Estimation and Western Blot Analysis**

The large right caudal lung lobe from each mouse was collected and homogenized in 300 μL RIPA buffer supplemented with a protease inhibitor cocktail (Thermo Fisher Scientific) using a PRO200 tissue homogenizer (PRO Scientific). Homogenates were vortexed every 30 minutes on ice (three cycles total) and centrifuged at 14,000 rpm for 20 minutes at 4°C. Clarified supernatants from lung homogenates were collected for protein estimation. Total protein concentration was measured using the Pierce™ Micro BCA Protein Assay Kit (Thermo Fisher Scientific, USA), following the manufacturer’s instructions.

For Western blotting, 25 μg of protein was resolved on 7.5% polyacrylamide gels and transferred onto nitrocellulose membranes using the Bio‑Rad Trans‑Blot Turbo system. Membranes were blocked for 1 hour at room temperature in 5% BSA in TBST (1× TBS with 0.05% Tween‑20), washed twice in TBST (5 minutes each), and incubated overnight at 4°C with primary antibodies (1:1000 dilution) targeting circadian clock proteins, adherens/tight junction markers, and signaling molecules. The list of primary and secondary antibodies used in the study is provided (**Table S1**). Following primary incubation, membranes were washed three times in TBST (10 minutes each) and incubated with appropriate HRP‑conjugated secondary antibodies for 1 hour at room temperature. After three additional washes, chemiluminescent substrate (Thermo Fisher Scientific) was applied, and signals were visualized using a Bio‑Rad ChemiDoc Imaging System. For detection of multiple targets on the same membrane, blots were stripped using commercial stripping buffer (Thermo Fisher Scientific), washed thoroughly, and reprobed with the appropriate primary antibody. Band intensities were quantified using ImageJ (NIH, USA), and protein expression levels were normalized to α‑tubulin as the loading control to calculate relative fold changes.

**References:**

1.	Srinivasan A, Giri A, Duraisamy SK, Alsup A, Castro M, Sundar IK. Acute HDM exposure shows time-of-day and sex-based differences in the severity of lung inflammation and circadian clock disruption. *J Allergy Clin Immunol Glob.* 2023;2(4):100155.

2.	Srinivasan A, Giri A, Duraisamy SK, Alsup A, Castro M, Sundar IK. Chronic HDM exposure shows time-of-day and sex-based differences in inflammatory response associated with lung circadian clock disruption. *iScience.* 2023;26(9):107580.
