## Supplemental Tables S1-S2 for "Circadian and Sex-Specific Responses to *Alternaria alternata* in Allergic Airway Inflammation"

**Table S1. Key Resources**

| **REAGENT or RESOURCE** | **SOURCE** | **IDENTIFIER** |
| --- | --- | --- |
| **Flow Cytometry Antibodies** | | |
| Purified anti-mouse CD16/32 antibody (clone 93) | BioLegend | Cat# 101302;  RRID:[AB_312801](http://antibodyregistry.org/AB_312801) |
| Ghost Dye Violet 510 | TonboBio | Cat# 13-0870-TT100 |
| Brilliant Violet 605 anti-mouse CD45 antibody (clone 30-F11) | BioLegend | Cat# 103155; RRID:[AB_2650656](http://antibodyregistry.org/AB_2650656) |
| PE/Cyanine7 anti-mouse CD3ε antibody (clone 145-2C11) | BioLegend | Cat# 100320; RRID:[AB_312685](http://antibodyregistry.org/AB_312685) |
| APC/Cyanine7 anti-mouse/human CD11b antibody (clone M1/70) | BioLegend | Cat# 101226; RRID:[AB_830642](http://antibodyregistry.org/AB_830642) |
| Brilliant Violet 650 anti-mouse I-A/I-E antibody (clone M5/114.15.2) | BioLegend | Cat# 107641; RRID:[AB_2565975](http://antibodyregistry.org/AB_2565975) |
| Brilliant Violet 785 anti-mouse CD11c antibody (clone N418) | BioLegend | Cat# 117336; RRID:[AB_2565268](http://antibodyregistry.org/AB_2565268) |
| Alexa Fluor 488 anti-mouse CD24 antibody (clone M1/69) | BioLegend | Cat# 101816;  RRID:[AB_493482](http://antibodyregistry.org/AB_493482) |
| PE anti-mouse CD193 (CCR3) antibody (clone J073E5) | BioLegend | Cat# 144506; RRID:[AB_2561534](http://antibodyregistry.org/AB_2561534) |
| Alexa Fluor 700 anti-mouse Ly-6G antibody (clone 1A8) | BD Biosciences | Cat# 561236; RRID:[AB_10611860](http://antibodyregistry.org/AB_10611860) |
| APC anti-mouse CD64 (FcγRI) antibody (clone X54-5/7.1) | BioLegend | Cat# 139306; RRID:[AB_11219391](http://antibodyregistry.org/AB_11219391) |
| PE-CF594 rat anti-mouse Siglec-F (clone E50-2440) | BD Biosciences | Cat# 562757; RRID:[AB_2687994](http://antibodyregistry.org/AB_2687994) |
| APC/Cyanine7 anti-mouse CD19 antibody (clone 6D5) | BioLegend | Cat# 115530;  RRID: AB_830707 |
| APC anti-mouse CD4 antibody (clone GK1.5) | BioLegend | Cat# 100412;  RRID: AB_312697 |
| Brilliant Violet 785 anti-mouse CD90.2 antibody (clone 30-H12) | BioLegend | Cat# 105331;  RRID: AB_2562900 |
| Brilliant Violet 711 anti-mouse T-bet antibody (clone 4B10) | BioLegend | Cat# 644820;  RRID: AB_2715766 |
| PE anti-mouse GATA-3 antibody (clone 16E10A23) | BioLegend | Cat# 653804;  RRID: AB_2562723 |
| PE-efluor^TM^ 610 Anti-mouse ROR gamma (t) antibody (clone B2D) | Invitrogen | Cat# 61-6981-82;  RRID: AB_2574650 |
| FITC anti-mouse NK1.1 antibody (clone PK136) | BioLegend | Cat# 108706;  RRID: AB_313393 |
| FITC anti-mouse TCR γ/δ antibody (clone GL3) | BioLegend | Cat# 118106;  RRID: AB_313830 |
| FITC anti-mouse Ly-6G antibody (clone 1A8) | BioLegend | Cat# 127606;  RRID: AB_1236494 |
| FITC anti-mouse CD5 antibody (clone 53-7.3) | BioLegend | Cat# 100606;  RRID: AB_312735 |
| FITC anti-mouse CD11b antibody (clone M1/70) | BioLegend | Cat# 101206;  RRID: AB_312789 |
| FITC anti-mouse CD11c antibody (clone N418) | BioLegend | Cat# 117306;  RRID: AB_313775 |
| FITC anti-mouse F4/80 antibody (clone BM8) | BioLegend | Cat# 123108;  RRID: AB_893502 |
| FITC anti-mouse FcεRIα antibody (clone MAR-1) | BioLegend | Cat# 134306;  RRID: AB_1626108 |
| **Western blot Antibodies** | | |
| CLOCK (D45B10) Rabbit Monoclonal Antibody | Cell Signaling  Technology | Cat# 5157S |
| BMAL1 (D2L7G) Rabbit Monoclonal Antibody | Cell Signaling  Technology | Cat# 14020 |
| E-Cadherin (24E10) Rabbit Monoclonal Antibody | Cell Signaling  Technology | Cat# 3195S |
| Phospho-Rev-erb alpha (Ser55/59) Antibody | Cell Signaling  Technology | Cat# 2129S |
| Rev-Erb-alpha (E1Y6D) Rabbit Monoclonal Antibody | Cell Signaling  Technology | Cat# 13418S |
| Phospho-Stat3 (Tyr705) (D3A7) Rabbit Monoclonal Antibody | Cell Signaling  Technology | Cat# 9145S |
| Stat3 (D3Z2G) Rabbit Monoclonal Antibody | Cell Signaling  Technology | Cat# 12640S |
| Phospho-Stat6 (Tyr641) (D8S9Y) Rabbit Monoclonal Antibody | Cell Signaling  Technology | Cat# 56554S |
| Stat6 (D3H4) Rabbit Monoclonal Antibody | Cell Signaling  Technology | Cat# 5397S |
| Occludin (E6B4R) Rabbit Monoclonal Antibody | Cell Signaling  Technology | Cat# 91131S |
| Phospho-GSK-3 alpha/beta (Ser21/9) (37F11) Rabbit Monoclonal Antibody | Cell Signaling  Technology | Cat# 9327S |
| GSK-3 alpha/beta (D75D3) Rabbit Monoclonal Antibody | Cell Signaling  Technology | Cat# 5676S |
| PER2 Rabbit pAb | Abclonal | Cat# A13168;  RRID: AB_2760019 |
| Phospho-STAT1 (Tyr701) Monoclonal Antibody (ST1P-11A5) | Invitrogen | Cat# 33-3400  RRID: AB_2533113 |
| STAT1 Monoclonal Antibody (STAT1-79) | Invitrogen | Cat# AHO0832  RRID: AB_2536317 |
| ZO-1 Monoclonal Antibody (ZO1-1A12) | Invitrogen | Cat# 33-9100;  RRID: AB_2533147 |
| Phospho-ERK1/ERK2 (Thr202, Tyr204) Polyclonal Antibody | Invitrogen | Cat# 36-8800;  RRID: AB_2533283 |
| ERK1/ERK2 Monoclonal Antibody (ERK-7D8) | Invitrogen | Cat# 13-6200;  RRID: AB_2533024 |
| Alpha Tubulin Monoclonal antibody | Proteintech | Cat# HRP-66031 |
| Goat anti-Rabbit IgG (H+L) Secondary Antibody, HRP | Invitrogen | Cat# 65-6120;  RRID: AB_2533967 |
| Goat anti-Mouse IgG (H+L) Secondary Antibody, HRP | Invitrogen | Cat# 31431;  RRID: AB_10960845 |
| **Chemicals and Reagents** | | |
| *Alternaria alternata* extract | Greer Laboratories | Cat# XPM1D3A25 |
| Stain Buffer (FBS) | BD Biosciences | Cat# 554656; RRID:[AB_2869006](http://antibodyregistry.org/AB_2869006) |
| **Biochemical Assays and Molecular Biology Kits** | | |
| OneComp eBeads Compensation Beads | ThermoFisher Scientific | Cat# 01-1111-42 |
| LEGENDplex Mouse Proinflammatory chemokine panel (13 plex) | BioLegend | Cat# 740007 |
| LEGENDplex Mouse T Helper Cytokine panel (12 plex) | BioLegend | Cat# 741044 |
| Mouse IL-33 Uncoated ELISA Kit | Invitrogen | Cat# 88-7333-88 |
| CyQUANT™ LDH Cytotoxicity Assay | Invitrogen | Cat# C20300 |
| Amplex™ Red Uric Acid/Uricase Assay Kit | Invitrogen | Cat# A22181 |
| RNeasy Mini kit | Qiagen | Cat# 74104 |
| High-Capacity cDNA Reverse Transcription Kit | Applied Biosystems | Cat# 4368814 |
| TaqMan™ Universal PCR Master Mix | Applied Biosystems | Cat# 4304437 |
| **Model Organism/Strain** | | |
| Mouse: C57BL/6NCr WT | NCI (Frederick, MD) | Strain Code: 556 |
| **Software** | | |
| FCS Express 7 | *De Novo* Software | [https://denovosoftware.com](https://denovosoftware.com/) |

**Table S2. Mouse gene-specific qRT-PCR primers used in this study**

| **Gene symbol** | **Forward Primer Sequence (5’-3’)** | **Reverse Primer Sequence (5’-3’)** |
| --- | --- | --- |
| ***Il4*** | **5’-GGTGTTCTTCGTTGCTGTGA-3’** | **5’-TCTCGAATGTACCAGGAGCC-3’** |
| ***Il6*** | **5’-GTCCTTCCTACCCCAATTTCCA -3’** | **5’-CGCACTAGGTTTGCCGAGTA -3’** |
| ***Il10*** | **5’-TGGGTTGCCAAGCCTTATCG -3’** | **5’-TTCAGCTTCTCACCCAGGGA -3’** |
| ***Il13*** | **5’-ATGTTGGTCAGGGAATCCAG-3’** | **5’-TGTGTCTCTCCCTCTGACCC-3’** |
| ***Il17*** | **5’-TTTAACTCCCTTGGCGCAAAA -3’** | **5’-CTTTCCCTCCGCATTGACAC -3’** |
| ***Tnfα*** | **5’-GATCGGTCCCCAAAGGGATG -3’** | **5’- TTTGCTACGACGTGGGCTAC-3’** |
| ***Ifnγ*** | **5’-CAGCAACAGCAAGGCGAAA -3’** | **5’-CTGGACCTGTGGGTTGTTGAC -3’** |
| ***Cxcl1*** | **5’-GGCTTCCTTATGTTCAAACAGGG -3’** | **5’-GCCGTTACTCGGGTAAATTACA -3’** |
| ***Nr1d1*** | **5’-GAGTCAGGGACTGGAAGCTG-3’** | **5’-AAGACATGACGACCCTGGAC-3’** |
| ***Cdh1*** | **5’-CAGTTCCGAGGTCTACACCTT -3’** | **5’-TGAATCGGGAGTCTTCCGAAAA -3’** |
| ***Ctnnb1*** | **5’-ATGGAGCCGGACAGAAAAGC -3’** | **5’-CTTGCCACTCAGGGAAGGA -3’** |
| ***Ocln*** | **5’-GCTGTGATGTGTGTTGACTG -3’** | **5’-CAAGGAGGTCCATCTGGCAG -3’** |
| ***Stat3*** | **5’-CACCTTGGATTGAGAGTCAAGAC -3’** | **5’-AGGAATCGGCTATATTGCTGGT -3’** |
| ***Stat6*** | **5’-CTCTGTGGGGCCTAATTTCCA -3’** | **5’-CATCTGAACCGACCAGGAACT -3’** |
| ***Rn18s*** | **5’-GTAACCCGTTGAACCCCATT-3’** | **5’-CCATCCAATCGGTAGTAGCG-3’** |
