## Supplemental Figures S1-S5 for "Circadian and Sex-Specific Responses to *Alternaria alternata* in Allergic Airway Inflammation"

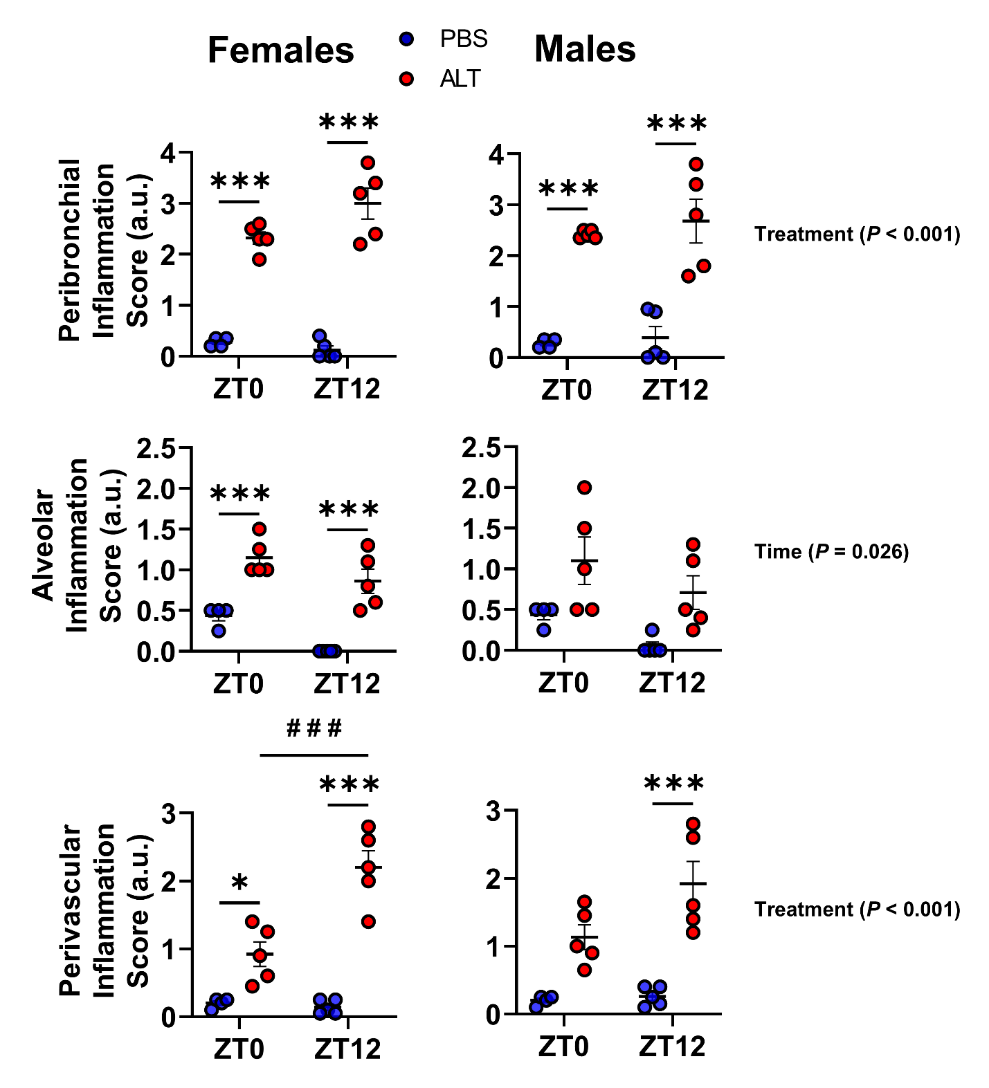

**Figure S1. Lung inflammation histopathology scores reveal region‑specific, time‑of‑day–dependent differences in ALT‑exposed female and male mice.** Hematoxylin and eosin–stained lung sections show varying inflammatory intensities in the peribronchial, perialveolar, and perivascular regions following PBS or ALT exposure in female and male mice. Graphs depict blinded semi‑quantitative inflammation scores as described in the Methods section. Data represent mean ± SEM (n = 4–5/group). Statistical analysis was performed using two‑way ANOVA followed by Tukey’s multiple comparison test. **P* < 0.05, *** *P* < 0.001, compared with the respective PBS control at ZT0 or ZT12; ^# # #^ *P* < 0.001, between ALT-exposed groups at ZT0 vs. ZT12. Interactions among Treatment x Sex x Time were analyzed using generalized linear modeling in R (see Supplemental Table S3).

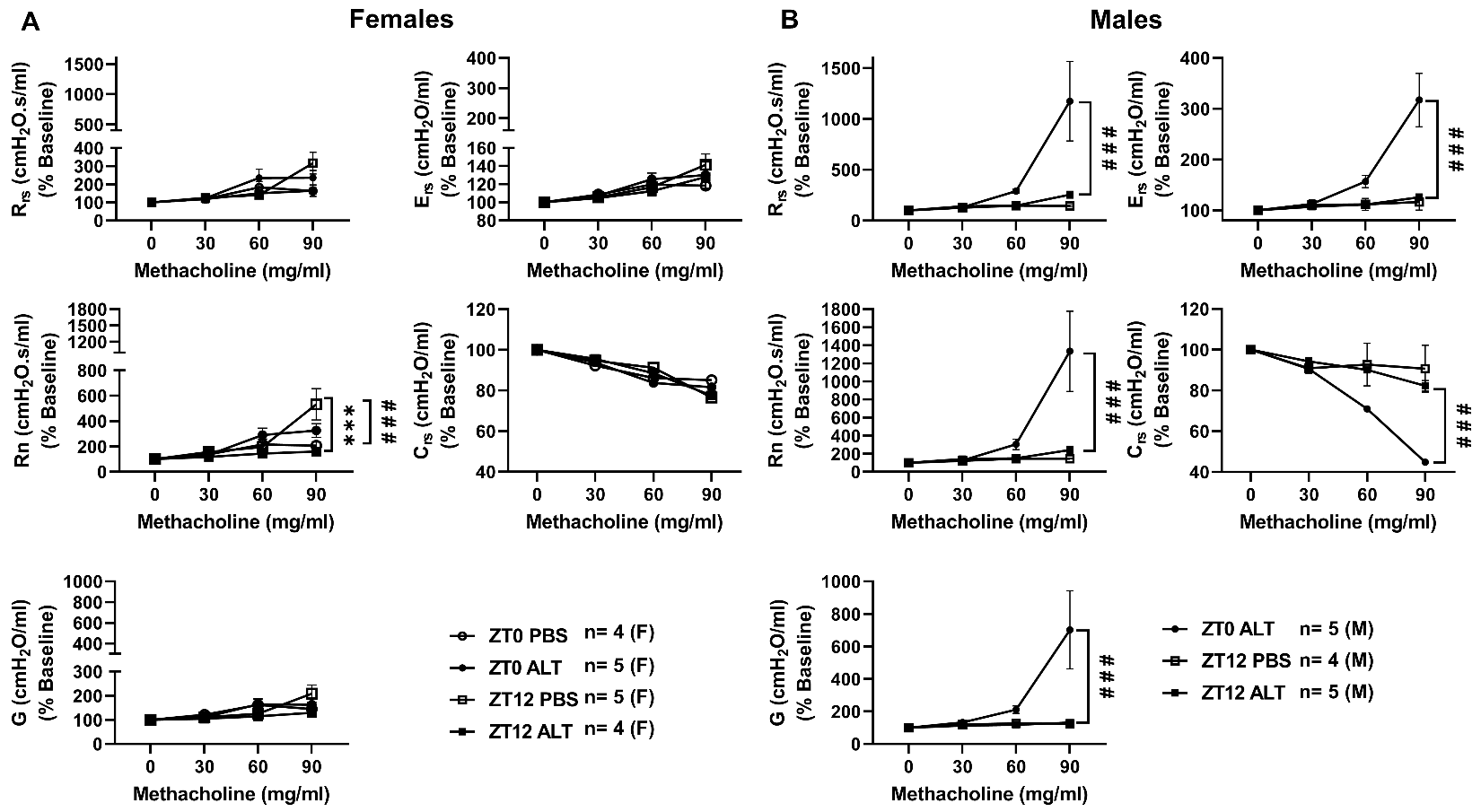

**Figure S2. *Alternaria alternata* (ALT) exposure produces differential lung function responses in female and male mice measured by the forced oscillation technique.** (A-B) C57BL/6 wild‑type female and male mice were administered ALT extract daily for four consecutive days, and lung function was assessed 24 hours after the final PBS or ALT exposure using the FlexiVent system. Forced oscillation technique parameters were normalized to baseline values at each concentration of aerosolized methacholine in PBS‑exposed mice at ZT0 and ZT12 (open circles and open squares) and ALT‑exposed mice at ZT0 and ZT12 (closed circles and closed squares). Values were normalized to individual baselines. Data represent mean ± SEM (n=4-5/group). Statistical analysis was performed using two‑way ANOVA followed by Tukey’s multiple comparison test. ^* * *^ *P* < 0.01, PBS ZT12 vs ALT ZT12 and ^# # #^*P* < 0.001, ALT ZT0 vs ALT ZT12.

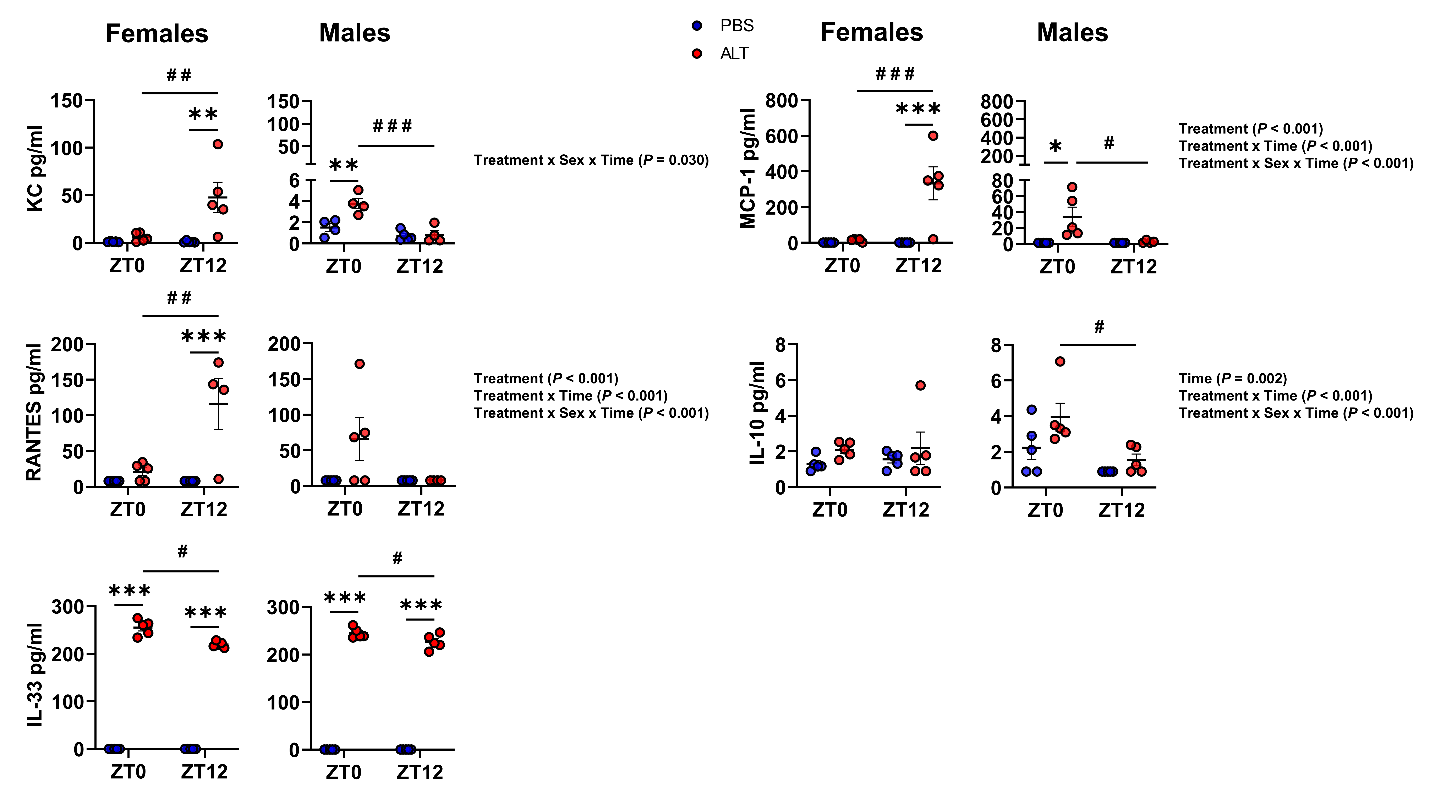

**Figure S3. Pro‑inflammatory mediators in BALF exhibit time‑of‑day and sex‑dependent differences in PBS‑ and ALT‑exposed female and male mice.** Wild‑type (WT) female and male mice were administered ALT extract daily for four consecutive days. Pro‑inflammatory mediators in BALF from PBS‑ and ALT‑exposed mice were quantified using a cytometric bead array (custom‑designed LEGENDplex Mouse Proinflammatory Cytokine/Chemokine panel). Data represent mean ± SEM (n = 4–5/group). Statistical analysis was performed using two‑way ANOVA followed by Tukey’s multiple comparison test. **P* < 0.05, ** *P* < 0.01, *** *P* < 0.001 compared with the respective PBS control at ZT0 or ZT12; ^#^ *P* < 0.05, ^# #^ *P* < 0.01, ^# # #^ *P* < 0.001 between ALT-exposed groups at ZT0 vs. ZT12. Interactions among Treatment x Sex x Time were analyzed using generalized linear modeling in R (see Supplemental Table S3).

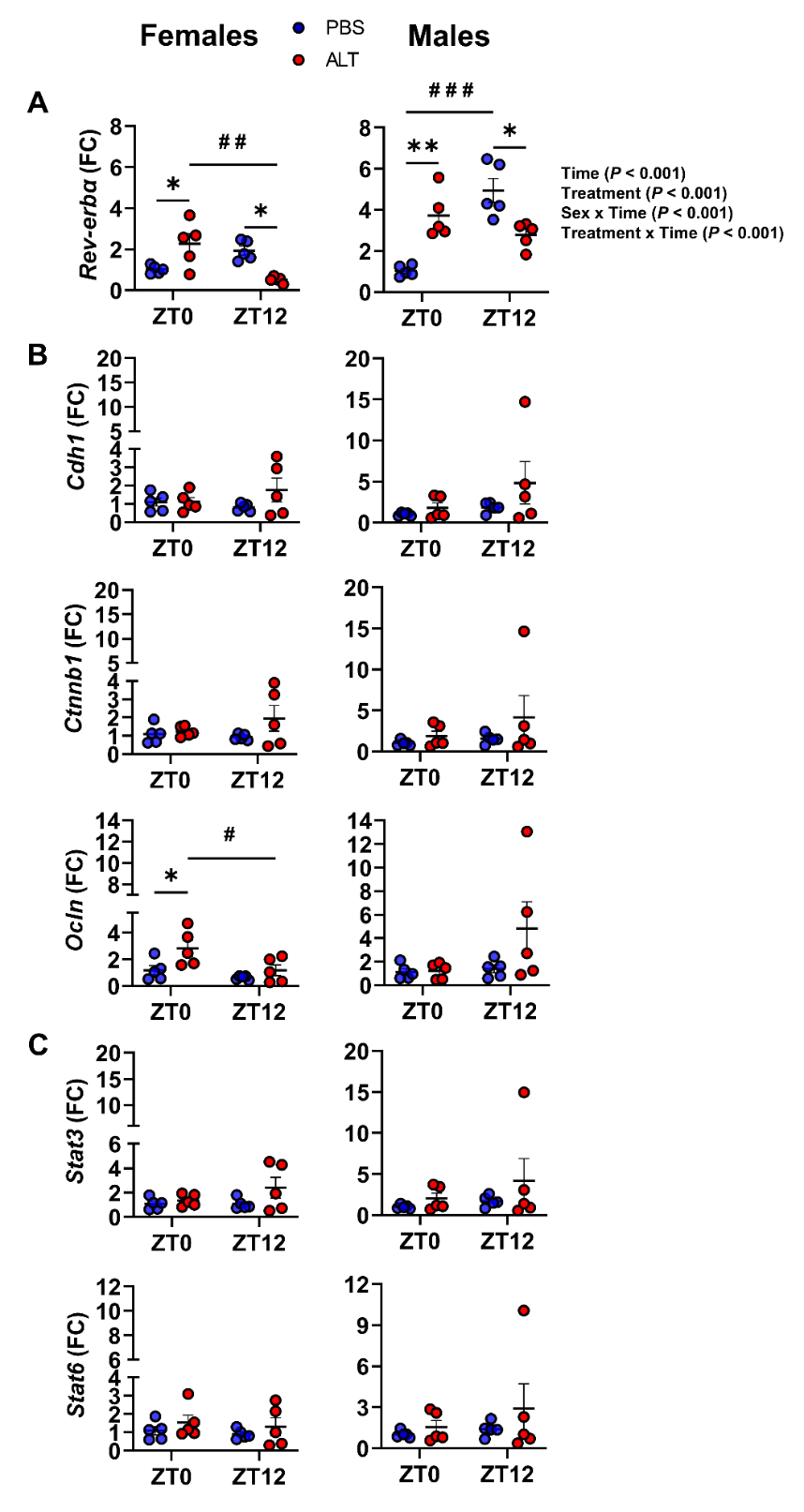

**Figure S4. *Alternaria alternata* (ALT) exposure elicits time‑of‑day– and sex‑dependent differences in gene expression of Rev‑erbα, epithelial barrier components, and transcription factors (STATs) in female and male mouse lungs.** Wild‑type (WT) female and male mice were administered ALT extract daily for four consecutive days. (A–C) Transcript levels of circadian clock (*Rev‑erbα*), epithelial barrier (*Cdh1, Ctnnb1, Ocln*), and transcription factor (*Stat3, Stat6*) genes in lung tissue were measured by quantitative real‑time qRT‑PCR and normalized to Rplp0 as the housekeeping gene. Data represent mean ± SEM (n = 4–5/group). Statistical analysis was performed using two‑way ANOVA followed by Tukey’s multiple comparison test. **P* < 0.05, ** *P* < 0.01 compared with the respective PBS control at ZT0 or ZT12; ^#^ *P* < 0.05, ^# #^ *P* < 0.01, ^# # #^ *P* < 0.001 between ALT-exposed groups at ZT0 vs. ZT12. Interactions among Treatment x Sex x Time were analyzed using generalized linear modeling in R (see Supplemental Table S3).

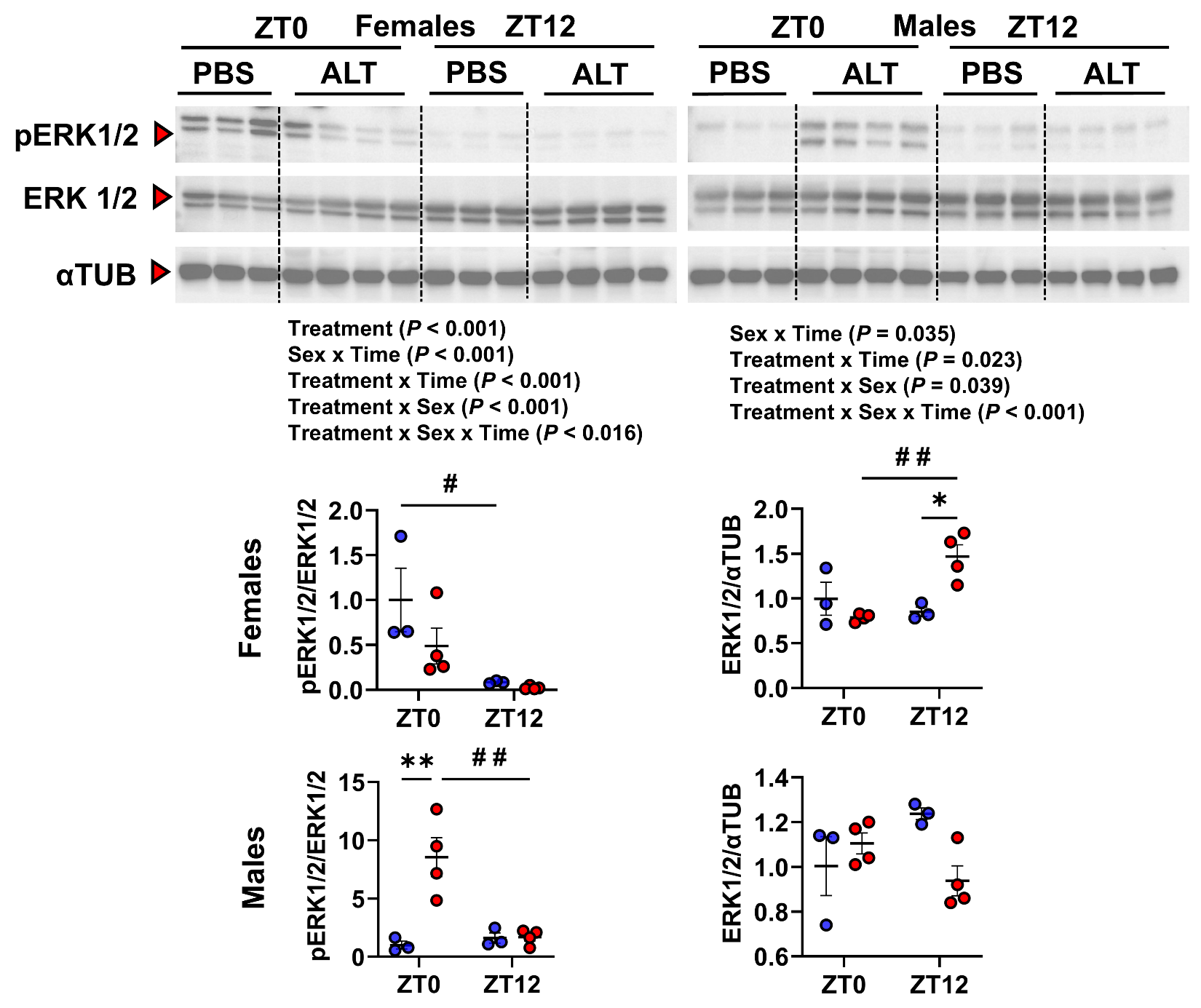

**Figure S5. *Alternaria alternata* (ALT)–induced ERK1/2 activation exhibits time‑of‑day– and sex‑dependent differences in female and male mouse lungs.** ALT exposure induced time‑of‑day–dependent changes in the protein expression of phosphorylated and total ERK1/2 in female and male mice. Representative Western blot images show alterations in phosphorylated and total ERK1/2 in PBS‑ and ALT‑treated groups at ZT0 and ZT12. α‑Tubulin was used as the loading control for densitometric analysis. Representative Western blot images were obtained from stripped and re‑probed membranes that were also used to detect other markers shown in the main figures (Figure 7A). Data represent mean ± SEM (n = 3–4/group). Statistical analysis was performed using two‑way ANOVA followed by Tukey’s multiple comparison test. **P* < 0.05, ***P* < 0.01 compared with the respective PBS control at ZT0 or ZT12; ^#^*P* < 0.05, ^# #^*P* < 0.01 between ALT-exposed groups at ZT0 vs. ZT12. Interactions among Treatment x Sex x Time were analyzed using generalized linear modeling in R (see Supplemental Table S3).
